# The Primate Cortical LFP Exhibits Multiple Spectral and Temporal Gradients and Widespread Task-Dependence During Visual Short-Term Memory

**DOI:** 10.1101/2024.01.29.577843

**Authors:** Steven J. Hoffman, Nicholas M. Dotson, Vinicius Lima, Charles M. Gray

## Abstract

Although cognitive functions are hypothesized to be mediated by synchronous neuronal interactions in multiple frequency bands among widely distributed cortical areas, we still lack a basic understanding of the distribution and task dependence of oscillatory activity across the cortical map. Here, we ask how the spectral and temporal properties of the local field potential (LFP) vary across the primate cerebral cortex, and how they are modulated during visual short-term memory. We measured the LFP from 55 cortical areas in two macaque monkeys while they performed a visual delayed match to sample task. Analysis of peak frequencies in the LFP power spectra reveals multiple discrete frequency bands between 3-80 Hz that differ between the two monkeys. The LFP power in each band, as well as the Sample Entropy, a measure of signal complexity, display distinct spatial gradients across the cortex, some of which correlate with reported spine counts in layer 3 pyramidal neurons. Cortical areas can be robustly decoded using a small number of spectral and temporal parameters, and significant task dependent increases and decreases in spectral power occur in all cortical areas. These findings reveal pronounced, widespread and spatially organized gradients in the spectral and temporal activity of cortical areas. Task-dependent changes in cortical activity are globally distributed, even for a simple cognitive task.

## Introduction

Extensive evidence, from studies in both humans and non-human primates, has established that cognitive tasks engage synchronized neuronal oscillations in multiple frequency bands among widely distributed cortical areas (Bressler et al., 1993; Tallon-Baudry et al., 2001; Jensen et al., 2002; Brovelli et al., 2004; Siegel et al., 2009; Liebe et al., 2012; Salazar et al., 2012; Dotson et al., 2014; Bastos et al., 2015; Lundqvist et al., 2016; Lobier et al., 2018; Rezayat et al., 2021; Vezoli et al., 2021). A number of functional roles have been proposed for these cortical rhythms, ranging from perceptual binding (Singer and Gray, 1995; Singer, 1999), working memory (Salazar et al., 2012; Miller et al., 2018; Rezayat et al., 2021) and attention (Fries 2005; 2015) to inhibitory gating mechanisms (Jensen and Mazaheri 2010; Hagan and Pesaran, 2022) and consciousness (Singer, 2001), often with the implicit assumption that the cortical rhythms of interest are expressed throughout the cortex. However, despite decades of research on the function and mechanisms of cortical oscillatory activity (Gray, 1994, Buzsáki and Draguhn 2004; Wang, 2010; Pesaran et al., 2018), it is still unclear how the various cortical rhythms are distributed across the cortical mantle.

Early electroencephalographic recordings of human brain activity revealed salient oscillations in the alpha (∼7-14 Hz) and beta (∼15-30 Hz) frequency bands (Berger 1929; Jasper and Andrews, 1936, 1938). Spatial mapping of the surface cortical EEG revealed a wide distribution of alpha activity throughout parietal, temporal, and occipital regions, as well as portions of the anterior frontal lobe and more localized beta activity in sensorimotor and premotor cortices (Jasper and Penfield 1949). Recent MEG and intracranial EEG studies have revealed distinct spectral features among the array of cortical areas (Keitel and Gross, 2016; Frauscher et al., 2018) and spatial gradients of peak frequency within the alpha and beta bands (Mahjoory et al., 2020). However, both approaches have significant limitations. Intracranial EEG recordings are limited in scope and the spatial resolution of MEG methods is hampered by the superposition of signals from multiple brain areas and the general dominance of the ongoing alpha rhythm (Keitel and Gross, 2016). Therefore, a high-resolution map of cortical oscillatory activity has remained elusive.

Elucidating the spatial and task-dependent organization of cortical oscillatory activity is an especially timely question, given recent discoveries of anatomical and functional gradients underlying the hierarchical organization of cortex (Goulas et al., 2018; Huntenburg et al., 2018; Hilgetag et al., 2019; Wang, 2020). Striking correlations have been established between gradients of dendritic spine counts (Elston, 2007; Elston et al., 20011), cell density (Collins et al., 2010; Beul et al., 2017), neurotransmitter receptor expression (Froudist-Walsh et al., 2021, 2023) and the anatomical hierarchy defined by interareal connections (Markov et al., 2014a,b; Chaudhury et al., 2015). Measures of the intrinsic time scale of neural activity across cortical areas also display a hierarchical organization consistent with the underlying anatomy (Hasson et al., 2008, 2015; Honey et al., 2012; Murray et al., 2014; Dotson et al., 2018; Gao et al., 2020; Spitmann et al., 2020). However, these functional studies do not include, or even discard, the oscillatory components of the signals under consideration. Consequently, a large gap exists in our understanding of how these anatomical and functional gradients are organized in relation to cortical oscillatory activity and its task dependence.

To address these questions, we developed and utilized a novel microdrive system to measure intracortical neural activity from a total of 55 identified cortical areas in two macaque monkeys that were engaged in a visual short-term memory task (Dotson et al., 2015, 2017, 2018). We measured both the power spectrum and the Sample Entropy (SampEn) (Richman and Moorman, 2000; Delgado-Bonal and Marshak, 2019) of the local field potential (LFP) to identify the predominant frequency bands of oscillatory activity, the relative and absolute amplitude of the signals in each band, and the temporal complexity of activity in each of the measured areas. These metrics displayed unique spatial gradients across the cortical areas, some of which were correlated with excitatory synaptic spine counts obtained from previous anatomical studies (Elston, 2007; Elston et al., 2011). Using the same metrics as features in a classifier, we found that cortical areas can be robustly decoded from each epoch of the task. All features and frequency bands contributed approximately equally to the classification performance. Task dependent changes in spectral power occurred in all cortical areas and nearly all frequency bands in both animals, revealing that even a simple cognitive task engages widespread areas of the cortical mantle (Gonzalez-Castillo et al., 2012).

## METHODS

Detailed descriptions of the experimental methods for behavioral training, neural recording, and preprocessing of the data have been described in two previous reports (Dotson et al., 2017; 2018). We provide brief descriptions of these methods here.

### Behavioral Task

Two female macaque monkeys were trained to perform an object-based delayed match to sample task (dMTS; MonkeyLogic software: Asaad and Eskandar, 2008a, 2008b). A schematic illustrating the time course of events in the task is shown in figure 1A. A trial began when the animal acquired and fixated on a small white fixation spot displayed on a grey background (presample period; fixation window = 3dva). After 500 ms for Monkey E, and 800 ms for Monkey L the fixation dot was replaced with one of five randomly selected sample images for 500 ms (size: 2.4 x 2.4 dva). The five daily images used were drawn from a larger pool of 92 images. During the sample period the monkey had to maintain its gaze within a 3 degree window encompassing the image. Next the sample image was extinguished and replaced with the fixation dot for a variable delay period (800-1200 ms Monkey E; 1000-1500 ms Monkey L). At the end of the delay period, the fixation target was extinguished, and the matching image and a non-matching image (one of the four other images) appeared at 5 degrees from the center of the screen. For Monkey E, the match and non-match images were always placed randomly in opposite hemifields along the horizontal axis. For Monkey L, the images were randomly aligned either vertically, horizontally or 45 degrees diagonally across from each other (see Fig. 1A). Finally, while the match images were visible, the monkey had to make a saccadic eye movement to the matching image and maintain fixation for a brief period (200 ms for Monkey E, and 500 ms for Monkey L). Correct trials were rewarded with a drop of juice.

**Figure 1.**
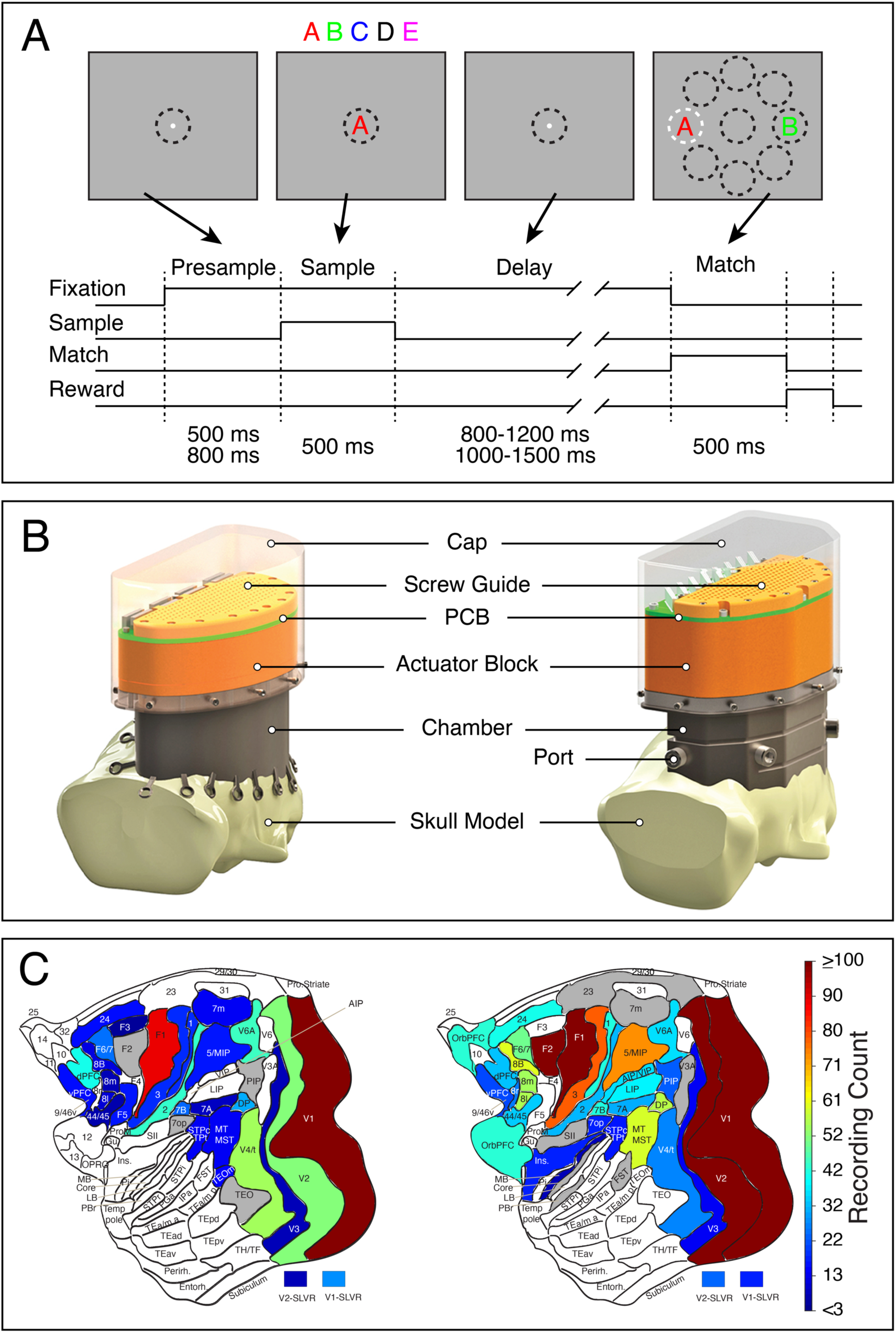
(A) Time course of events in the delayed match-to-sample task. The upper frames illustrate the sequence of images viewed by the monkeys during the task. The dashed circles represent the windows for monitoring eye position. These were not visible to the animal. The sample and match stimuli are symbolized by the letters A-E. The white dashed circle in the right plot indicates a correct match. The lower plot illustrates the timeline of task events. The presample duration was 500 ms for monkey E and 800 ms for monkey L. The duration of the delay period was 800-1200 ms for monkey E and 1000-1500 ms for monkey L. Match stimuli were randomly presented on opposite sides of the horizontal for monkey E and on opposite sides of axes lying at 0, 45 and 90 degrees for monkey L. (B) Design drawings of the chamber and semi-chronic microdrive systems used for recording neuronal activity. The left and right designs were used on Monkeys E and L, respectively. Adapted from Dotson et al. (2017). (C) Flatmaps of the recording counts for each cortical area/group (following the nomenclature of Markov et al, 2014a) in monkeys E (left) and L (right). Areas with less than 3 recordings are shaded grey and were not included in the analyses. Areal groupings and recording counts are listed in Table 1.

**Table 1.** Recording counts for all areas and areal groupings in Monkeys E and L. Area and areal group names are given in the first and second columns, respectively. Columns labeled as “Flatmaps” show the recording counts reported in flatmaps and rank ordered plots of SC, PA and SampEn. Columns labeled as “Decoding” show the area/groups that are included in the classification analysis.

| Areas | Area/Group | Monkey E | Monkey E | Monkey L | Monkey L |
| --- | --- | --- | --- | --- | --- |
|  |  | Flatmaps | Decoding | Flatmaps | Decoding |
| OPRO, 11,12,13,14,32 | OrbPFC |  |  | 42 | 42 |
| 24c, 24d | a24 | 14 |  | 40 | 40 |
| 9/46v, 46v | vPFC | 14 |  | 20 | 20 |
| 9/46d, 9, 46d | dPFC | 40 | 40 | 32 | 32 |
| 8B | 8B | 10 |  | 60 | 60 |
| 8L | 8L | 9 |  | 55 | 55 |
| 8M | 8M | 6 |  | 54 | 54 |
| 8r | 8r |  |  | 25 | 25 |
| 44, 45A, 45B | a44/45 | 5 |  | 33 | 33 |
| F1 | F1 | 90 | 90 | 102 | 102 |
| F2 | F2 |  |  | 120 | 120 |
| F3 | F3 | 3 |  |  |  |
| F5 | F5 | 15 |  |  |  |
| F6, F7 | F6/F7 | 24 | 24 | 49 | 49 |
| 1 | 1 | 17 |  | 41 | 41 |
| 2 | 2 | 43 | 43 | 31 | 31 |
| 3 | 3 | 17 |  | 79 | 79 |
| Insula | Ins |  |  | 16 |  |
| MB | MB |  |  | 6 |  |
| 7op | 7op |  |  | 16 |  |
| 7m | 7m | 12 |  |  |  |
| a5, MIP | a5/MIP | 15 |  | 74 | 74 |
| PIP | PIP |  |  | 22 | 22 |
| 7a | 7a | 7 |  | 30 | 30 |
| STPc, TPt | STPc/TPt | 12 |  | 15 |  |
| AIP, VIP | AIP/VIP |  |  | 35 | 35 |
| 7b | 7b | 23 | 23 | 41 | 41 |
| LIP | LIP |  |  | 37 | 37 |
| DP | DP | 30 | 30 | 56 | 56 |
| V6A | V6A | 44 | 44 | 35 | 35 |
| MT, MST | MT/MST | 12 |  | 59 | 59 |
| TEOm | TEOm | 13 |  |  |  |
| V4, V4t | V4/t | 54 | 54 | 26 | 26 |
| V3 | V3 | 6 |  | 12 |  |
| V2SLVR | V2SLVR | 6 |  | 23 | 23 |
| V2 | V2 | 51 | 51 | 117 | 117 |
| V1SLVR | V1SLVR | 27 | 27 | 16 |  |
| V1 | V1 | 169 | 169 | 225 | 225 |
| Recording count |  | **<u>788</u>** | **<u>526</u>** | **<u>1644</u>** | **<u>1563</u>** |
| Area/groups |  | 29 | 11 | 34 | 28 |

### Electrophysiological recording

Broadband recordings (0.1 Hz – 9 kHz, sampled at 32 kHz) were made across the breadth of a cerebral hemisphere in the two monkeys as described in Dotson et al., 2017. Briefly, a hemisphere-wide, large-scale microdrive was implanted in two female macaque monkeys with the capability of simultaneously recording from up to 256 independently moveable micro-electrodes (inter-electrode spacing = 2.5 mm) (Figure 1B). Neural activity was sampled from an overlapping set of 62 cortical areas over the course of 6 and 9 months for Monkeys E and L, respectively (Figure 1C). The broadband signal was bandpass filtered at 1-250 Hz (4th order Butterworth) and resampled at 1kHz in order to obtain the LFP. Anatomical designations (using the nomenclature of Markov et al., 2014a) were achieved through reconstruction of each electrode!s track through histological sections (Dotson et al., 2017). Only recordings with neural unit activity that exceeded a minimum mean firing rate of 1 Hz were used for further analysis. Spike waveforms were extracted by detecting local minima in the highpass signal (500 Hz - 9 kHz) that exceeded 5 SDs of the noise level (Yen et al., 2007; Salazar et al., 2012; Dotson et al., 2014, 2018). To avoid the over-sampling of activity, only LFP signals where the electrode position differed from the previous day of recording by >250 µm in depth were used in this analysis.

### Areal grouping

Data from adjacent cortical areas with few recordings and similar functional properties were pooled to form small groups of areas. Recordings in visual areas V1 and V2 with short-latency visual responses in the unit activity (SLVR) were analyzed separately (i.e., V1-SLVR, V2-SLVR). The SLVR designation was assigned if the unit activity displayed a significant change in firing rate within 50-100 ms following the sample presentation. After these pooling procedures, there were a total of 29 and 34 different cortical areas or small groups of areas in monkeys E and L, respectively (Table 1).

### Identification of spectral bands

For each recording the power spectral density, averaged across all correct trials in a session, was calculated from 1 – 100 Hz (Field Trip toolbox; Oostenveld et al., 2011). A multi-taper smoothing window of 1Hz was used. In Monkey E residual 60 Hz line noise was removed with a notch filter. Spectra were calculated over a time window of 2100 ms encompassing the task period from presample until match onset. Due to the variable length of the delay period, trials that were less than 2100 ms in length were zero padded in order to achieve this length. Longer trials were truncated to fit the 2100 ms time window. The frequency resolution of the power spectra was 0.5 Hz.

We used an empirical method to designate the frequency band limits for further analysis. We applied a peak detection algorithm (findpeaks.m in Matlab) to the normalized power spectra from 4-100 Hz, in both linear and semi-log coordinates, to identify narrow band oscillatory activity. Local maxima <4Hz were excluded, due to artifactual peaks introduced by the multi-taper smoothing filter. In order to limit small, spurious peaks, a peak prominence criterion (as defined by the Matlab findpeaks.m function) was used. The prominence threshold was set at 1% of the maximum prominence found across all spectra. Values exceeding the prominence threshold were used to generate a histogram of peak frequencies across all recordings in each animal. The histograms were fit with a probability density function (PDF) using a mixture of Gaussians (Fitdist.m function in Matlab). Frequency band limits were designated as the local minima in the PDFs. This analysis yielded nearly identical band limits when applied to the linear and semi-log spectra, with deviations in frequency bands being less than 1.5 Hz. Results from the two approaches were combined by averaging the local minima designations from the two methods.

### Classifier features

We first computed the average power spectral density (1 Hz multi-taper smoothing window) across all trials for each of five 400 ms epochs of the task (see Figure 2B) on each recording (Field Trip toolbox; Oostenveld et al., 2011). Using the band limits defined for each monkey, we measured the spectral content (SC) and the peak amplitude (PA) of the spectrum in each frequency band and epoch of the task for the entire data set in each animal. We defined SC as the percentage of power within each frequency band relative to the entire spectrum (0-80 Hz) and PA as the largest value of the average spectrum in each band.

**Figure 2.**
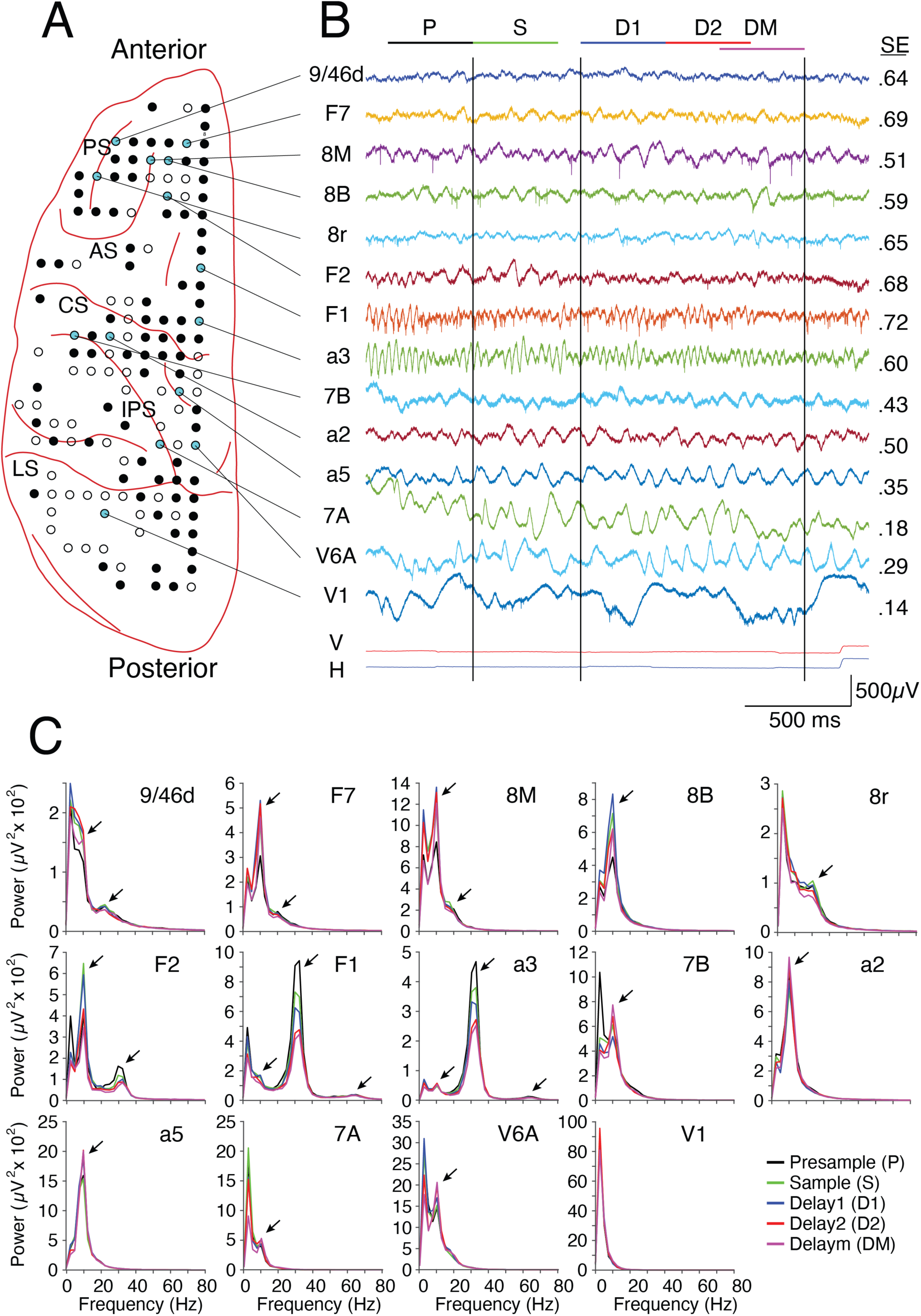
Spectral properties of LFP signals vary markedly across cortex. (A) Schematic of the recording sites in monkey L. The outline of the left hemisphere and major cortical sulci, drawn from a photograph, are shown in red. Each circle shows the entry location of an electrode that recorded neural unit activity at some point during the 9 months of the experiment. Filled circles indicate electrode locations that recorded neural activity in this session (n=94). Cyan filled circles mark the electrode locations that correspond to the signals shown in B. (B) Broadband (0.1Hz-9kHz) raw data recorded on a single trial from 14 separate cortical areas. The areal name (left of each trace) follows the nomenclature of Markov et al. (2014a). The bottom two traces show the vertical and horizontal components of the eye position signal. The black vertical lines, from left to right, mark the onset and offset of the sample stimulus and the onset of the match stimulus, respectively. The colored horizontal bars at the top indicate the time and duration (400 ms) of the Presample (P), Sample (S), Delay1 (D1), Delay2 (D2) and DelayM (DM) epochs, respectively. A saccadic eye movement, occurring ∼200 ms following the onset of the match, indicates the monkey!s behavioral choice. The mean Sample Entropy (SampEn) across trials is plotted to the right of each data trace. (C) LFP power spectra (0-80 Hz) for each area indicated in A and B, averaged across all correct trials for each of the 5 epochs (Presample: black; Sample: green; Delay1: blue; Delay2: red; Delaym: magenta). Clear differences in LFP power and its task dependence are apparent between different areas of cortex. Black arrows mark notable local peaks or shoulders in the power spectra. Peaks occurring below 4 Hz are due to the absence of a DC component in the filtered signals. Abbreviations: PS - Principal Sulcus; AS – Arcuate Sulcus; CS – Central Sulcus; IPS – Intraparietal Sulcus; LS – Lunate Sulcus.

To quantify the complexity, or degree of randomness, of the LFP, we calculated the Sample Entropy (SampEn(m,r,N)) on each task epoch of each trial for each recording (Richman and Moorman 2000). We used the average value of SampEn across trials for each epoch as a feature. SampEn is an event counting statistic, derived from Approximate Entropy (ApEn) (Pincus, 1991), which quantifies the persistence or regularity of similar patterns in a signal. It is defined as “*the negative natural logarithm of the conditional probability that two sequences similar for m points remain similar at the next point, where self-matches are not included in calculating the probability*” (Richman and Moorman, 2000; Delgado-Bonal and Marshak, 2019). It reduces the bias associated with ApEn and is largely independent of the record length of the data.

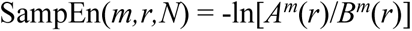

where

*B^m^*(*r*) is the probability that two sequences are similar for *m* points (possibles)

*A^m^*(r) is the probability that two sequences are similar for *m+1* points (matches)

*m* is the length of the sequences being compared

*r* is the threshold for accepted matches, expressed as a fraction of the standard deviation (σ) of the time series

*N* is the length of the time series being evaluated

We used values of *m*=2 and *r*=0.2σ. *N*=400 data points, which is the length of each task epoch. Each data epoch of length N was z-score normalized before computing SampEn.

Together these analyses resulted in 9 features in each of 5 epochs in monkey E (SC and PA in bands 1-4, and SampEn) and 11 features in each of 5 epochs in monkey L (SC and PA in bands 1-5, and SampEn).

### Machine learning classifier

Each of the features were included in a cubic Support Vector Machine (SVM) classifier. Cortical areas with small numbers of recordings were grouped with those in adjacent areas with similar functional properties (Table 1). To avoid bias (in both the training and validation phases) towards areas with substantially more recordings, an equal number of recordings (n=20) from each area/group were randomly drawn from the larger pool of data. The classifier was trained and validated using k-fold cross validation (k=5, 4/5^th^ training, 1/5^th^ cross validation). The training-validation process was iteratively repeated 1000 times, with each iteration using a different set of 20 randomly selected recordings for an area/group drawn from the data pool (Table 1). The theoretical level of chance classification is 1/# areas (1/11 Monkey E, 1/28 Monkey L). In order to further assess classification accuracy and avoid biases arising from the use of the theoretical chance level (Combrisson and Jerbi 2015), the mean and 99% confidence interval for chance classification was calculated from a permutation distribution. This distribution was created by randomly shuffling areal/group labels before the classification process, then repeating over 1000 iterations.

## RESULTS

We recorded broadband neuronal activity from a total of 55 cortical areas in two rhesus macaque monkeys (33 areas in Monkey E and 50 areas in Monkey L) while they performed a feature-based delayed match-to-sample (dMTS) task and an interleaved visual fixation task (Figure 1A). The dMTS task required the monkeys to remember a centrally presented sample image (1 of 5 possible images) for a minimum of 800 ms (800-1200 ms in Monkey E; 1000-1500 ms in Monkey L), before making a choice between a matching and a non-matching image. Details of the recording methods, the behavioral task, reconstruction of the recording sites and analysis of the task dependence of unit activity have been reported previously (Dotson et al., 2017, 2018). Here we focus on the spatial and spectro-temporal organization of the LFP and its task dependence. LFP signals were selected for analysis only if there was detectable unit activity recorded on the same electrode, and if the electrode position had changed by more than 250 µm from the previous session of recording, yielding a unique recording site. LFP signals that were noisy or had frequent artifacts were discarded, following the methods described in Salazar et al. (2012). All analyses were performed on the correct trials of the dMTS task, obtained from 25 sessions in Monkey E and 61 sessions in Monkey L (minimum 500 correct trials and >75% correct performance on each session). Simultaneous recordings were made from up to 21 and 37 different cortical areas in Monkeys E and L, respectively. The cortical area of the recording locations and sample sizes for each animal are shown in Figure 1C and Table 1. Data from sparsely sampled and adjacent cortical areas with similar functional properties were merged into small groups. This resulted in a total of 788 recordings from 33 areas merged into 29 area/groups in Monkey E and a total of 1644 recordings from 50 areas merged into 34 area/groups in Monkey L (Table 1). For simplicity we use the term “areas” throughout the text when describing our findings for cortical areas and small groups of areas.

### Spatial Organization of the LFP

At the outset of these experiments, it was apparent that the spectral and temporal properties of the LFP varied systematically across the cortex in a task dependent manner. Figure 2 shows a representative example of the raw data, and corresponding LFP power spectra, sampled from a subset of the channels on a single session in monkey L. Prefrontal (9/46d), frontal eye field (8M, 8B, 8r), and premotor (F7, F2) areas tended to display desynchronized fluctuations of low amplitude interspersed with brief intervals of periodic activity in the range of 6-14 Hz. Primary motor and somatosensory areas, F1 and a3, exhibited pronounced oscillations in the 25-35 Hz range, that occurred with lower amplitude in premotor areas. Anterior (a2, a5, 7B) and posterior (7A, V6A) parietal areas exhibited higher amplitude fluctuations with salient oscillations in the 6-14 Hz range. Primary visual cortex (V1) was dominated by high amplitude fluctuations at low frequencies. Each of these aspects were clearly apparent in the corresponding power spectra (Figure 2C), which were computed from five separate 400 ms epochs of the task (presample (P), sample (S), early delay (D1), late delay (D2), match locked delay (DM), Figure 2B). There were notable peaks in the spectra which often varied between task epochs. Some spectra showed a sharp singular peak centered around 10 Hz (Figure 2C, areas 8B, 7B, a2, a5, 7A, V6A), while others displayed multiple peaks around 10 Hz and 30 Hz (Figure 2C, areas F2, F1, a3), and in some cases small peaks near 65 Hz (Figure 2C, areas F1, a3). Smaller peaks and shoulders were also apparent near 20 Hz (Figure 2C, areas 9/46d, F7, 8M, 8r). Other recordings, most notably in early visual cortex, displayed a steep fall off in power, with very low amplitudes at higher frequencies (Figure 2C, area V1).

To determine the appropriate frequency bands for subsequent analysis, we identified local peaks in the average power spectra, in both linear and semi-log coordinates, computed across all correct trials over the full trial length for all recordings in both animals (Figures S1 and S2). This analysis yielded a histogram of peak frequencies, and their corresponding areas of origin, spanning the full data set in each monkey (Figure 3). The histograms derived from the linear and semi-log spectra were nearly indistinguishable and had no effect on the selection of frequency bands (Figure S3). Frequencies below 4 Hz were excluded to avoid the detection of spurious peaks due to filtering. We fit each histogram with a probability density function (PDF) and found the local minima in the distributions. This resulted in 4 frequency bands in monkey E (0-8 Hz, 8-21 Hz, 21-32 Hz, 32-80 Hz) and 5 bands in monkey L (0-6 Hz, 6-14 Hz, 14-26 Hz, 26-42 Hz, 42-80 Hz). These bands did not change after removing the data from all areas that were not present in both animals (V1SL in Monkey E and OrbPFC, 8r, Ins, AIP/VIP, LIP in Monkey L).

**Figure 3.**
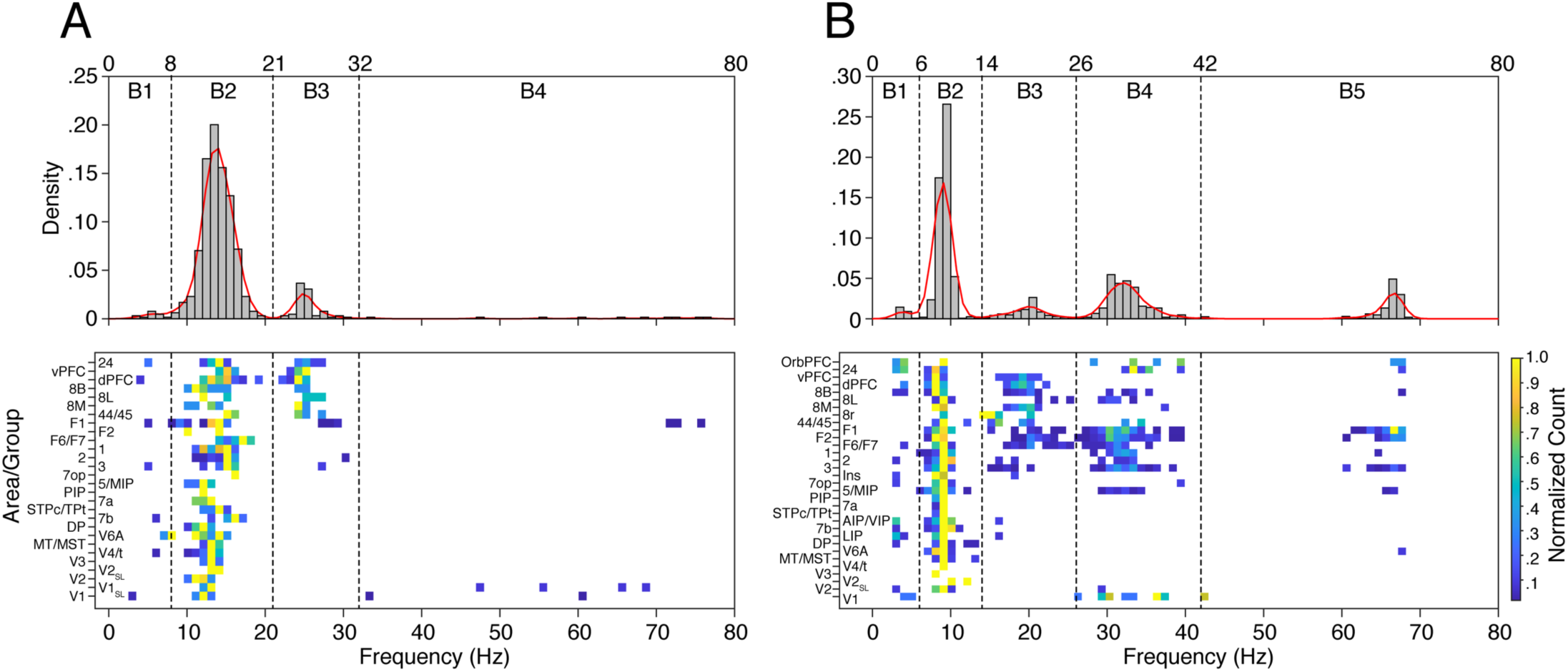
Spectral peaks in the LFP occur in distinct frequency bands that vary between animals. Distributions of peak frequencies obtained from semi-log power spectra on all channels and sessions in monkeys E (A) and L (B). The top plots in A and B show the cumulative histograms of all peak frequencies (4-80 Hz) that exceeded the peak prominence threshold (see Figures S1 and S2). The continuous red lines show the probability density function (PDF) computed with a mixture of Gaussians fit to each distribution. The local minima in each PDF define the boundaries between selected frequency bands for each animal (dashed vertical lines). Four bands (B1-B4) were chosen for Monkey E and five bands (B1-B5) were chosen for Monkey L. The lower plots show the normalized counts of peak frequencies for each cortical area/group. This revealed a rough spatial organization of peak frequencies across the sampled cortical areas.

These data revealed a notable difference in the frequency distribution of narrow-band oscillations between the two monkeys. Low frequency spectral peaks (band-1) were sparsely distributed in both monkeys. Spectral peaks in band-2, centered at 14 Hz in monkey E and 10 Hz in monkey L, were widely distributed and present in nearly all areas of cortex. Peak frequencies in band-3 were centered at 25 Hz in monkey E and 21 Hz in monkey L and were frontally distributed. There were more obvious differences between the animals at higher frequencies. Spectral peaks in band-4 were very sparse in monkey E, while monkey L showed pronounced narrow-band components in two bands (band-4, band-5) that were concentrated in somatomotor regions and some areas of the prefrontal and parietal cortices. The spectral peaks in band-5 of monkey L were notable in that, with one exception, they always co-occurred with and had twice the frequency of the peaks in band-4. This suggested the presence of a higher frequency harmonic known to result from the non-sinusoidal nature of oscillations in LFP, EEG and MEG signals (Aru et al., 2015; Lozano-Soldevilla et al., 2016; Gerber et al., 2016; Cole et al., 2017; Schaworonkow and Nikulin, 2019). Detailed analysis of the band-5 component, in both the frequency and time domains (see Frequency Harmonic Analysis in Supplementary Materials), confirmed this conjecture (Figures S4, S5, S6, S7). Some high frequency components were also sparsely present in early visual cortex in both animals, consistent with the occurrence of induced gamma-band activity in response to the sample stimuli (Friedman-Hill et al., 2000). Many areas displayed a unimodal distribution, while other areas showed bimodal or even multimodal distributions of spectral peaks. This was most evident in monkey L. These results reveal distinct differences between animals in the frequency composition of narrow band oscillations in the cortical LFP.

To determine if cortical areas can be distinguished by the spectral properties of their LFP, we extracted two values from the spectra computed on each epoch: the percentage of power in each frequency band relative to the entire spectrum (0-80 Hz), which we refer to as the spectral content (SC), and the peak amplitude (PA) of the power in each band. The PA was distinct from SC because it reflected the absolute amplitude of the signals, which varied widely across cortical areas, and was not bounded within a range of 0-100% (Figure 2C). To assess the spatial organization of spectral power across the cortex, we plotted the mean value of SC in the presample epoch (across sessions) on cortical flatmaps (adopted from Markov et al., 2014a) for each frequency band (Figure 4). The corresponding rank ordered box plot of SC (sorted on the median value) is displayed to the right of each flatmap to visualize the distribution of values across cortical areas. These maps revealed striking spatial gradients of SC that differ between frequency bands and display similarities as well as differences between the two animals. In both monkeys there was a clear posterior to anterior gradient of SC in band-1, with the greatest concentration of power occurring in visual cortices. Band-2 displayed an anterior shift in the distribution with relative power radiating out from more central/parietal regions. In the higher frequency bands (3 and 4 in Monkey E (Figure 4A), 3-5 in Monkey L (Figure 4B)) there was an anterior shift in the distribution of SC with increasing frequency. A notable difference between animals was also present in the higher frequency bands. Monkey L displayed a distinct focal distribution of high amplitude (band-4) in somatomotor areas (3, F1 and F2) with a declining gradient into premotor, prefrontal and anterior parietal areas that was not present in monkey E. Similar plots of PA are shown for both monkeys in figure 5. These data reveal pronounced amplitude differences across areas within each band as well as striking differences in the LFP amplitude between different frequency bands. The spatial gradients are also apparent, but more variable than those shown by the normalized measure of SC in Figure 4.

**Figure 4.**
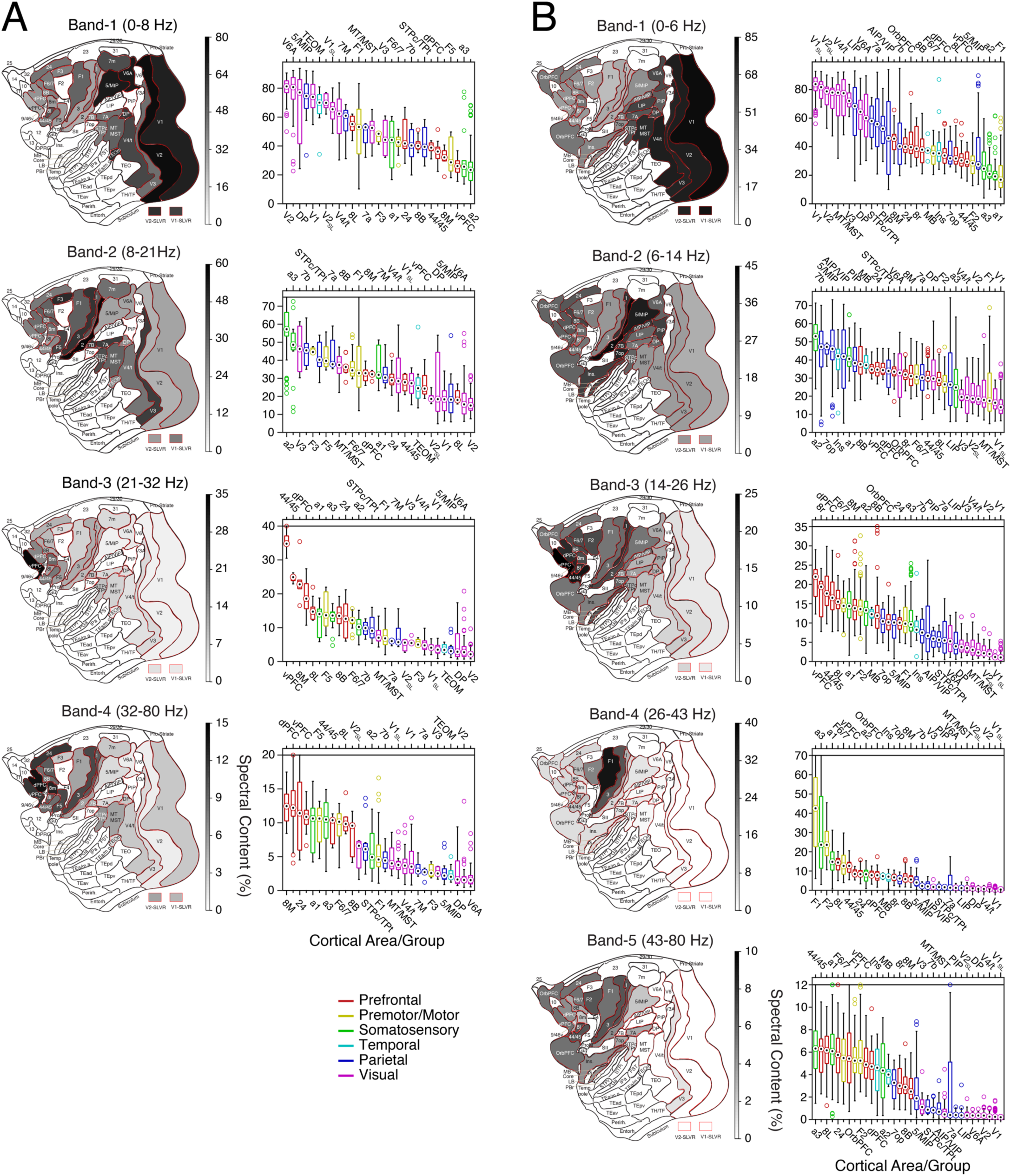
Spectral content displays anatomical gradients that differ across frequency bands in both monkeys (A: monkey E; B: monkey L). The left columns in A and B show cortical flatmaps of the mean spectral content in each area/group, across all sessions, during the presample epoch for each frequency band. Areal boundaries and nomenclature follow that of Markov et al (2014a). V1-SLVR and V2-SLVR refer to the subset of recordings in areas V1 and V2, respectively, that displayed short latency responses of spiking activity to one or more of the sample stimuli presented on each session (Dotson et al., 2018). The right columns in A and B show the corresponding distributions of spectral content across all sessions, ranked by the median values. The circle within each box shows the median, the box displays the interquartile range, the whiskers show the 5^th^ and 95^th^ percentiles, and the open circles show outliers. Areal labels are displayed at the bottom and top of each plot. The data are color-coded by cortical region (Prefrontal: red; Premotor/Motor: yellow; Somatosensory: green; Temporal: cyan; Parietal: blue; Visual: magenta).

**Figure 5.**
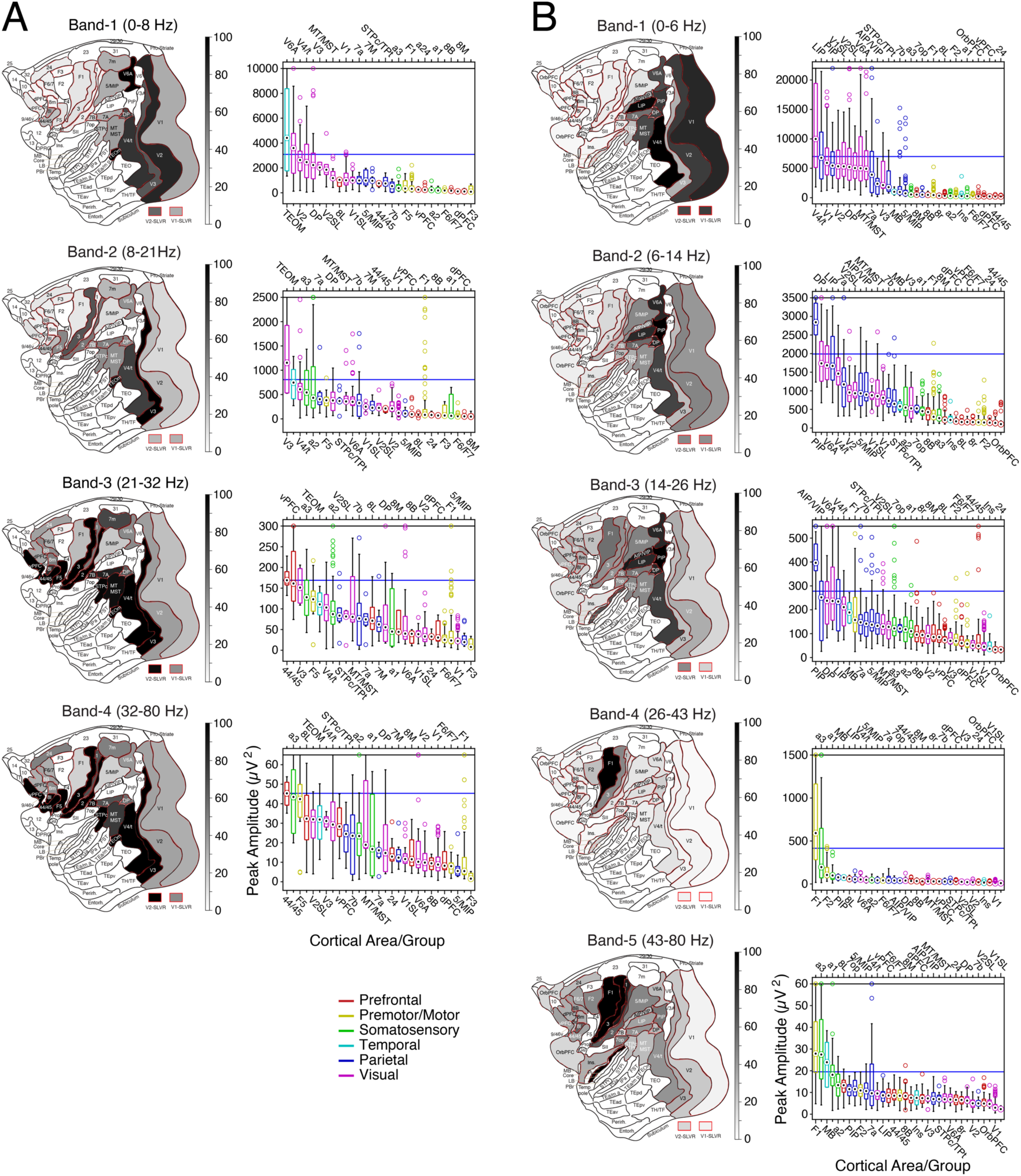
Flatmaps and rank ordered box plots of the peak amplitude (PA) obtained from the power spectra in each frequency band in Monkey E (A) and Monkey L (B) during the presample epoch of the task. Plotting conventions are the same as figure 4. Because of outliers and nonlinearities in the rank order plots, the flatmaps are scaled as a percentage of a threshold value shown by the blue line in each rank ordered plot.

The LFP also displayed marked variation in temporal structure across different areas of cortex (Figure 2B). To quantify this, we computed the Sample Entropy (SampEn) of each LFP time series (Richman and Moorman, 2000; Delgado-Bonal and Marshak, 2019) (see Methods). SampEn is an event counting statistic, derived from Approximate Entropy (ApEn) (Pincus, 1991), which quantifies the persistence or regularity of similar patterns in a signal. It reflects the degree of a signal!s randomness or complexity. A low value of SampEn indicates more self-similarity in a time series of LFP data. For each channel on each session, we calculated the average SampEn across trials on each epoch of the task. Examples of the mean SampEn values, averaged across epochs, are plotted to the right of the data traces in Figure 2B. SampEn was lower in posterior occipital and parietal areas and increased in somatomotor and prefrontal areas. This effect was consistent with the data as a whole. Flatmaps and corresponding rank ordered box plots of SampEn (presample epoch) for the full data set are shown for both monkeys in Figure 6. This revealed a clear spatial gradient of SampEn from occipital to prefrontal regions of the cortex.

**Figure 6.**
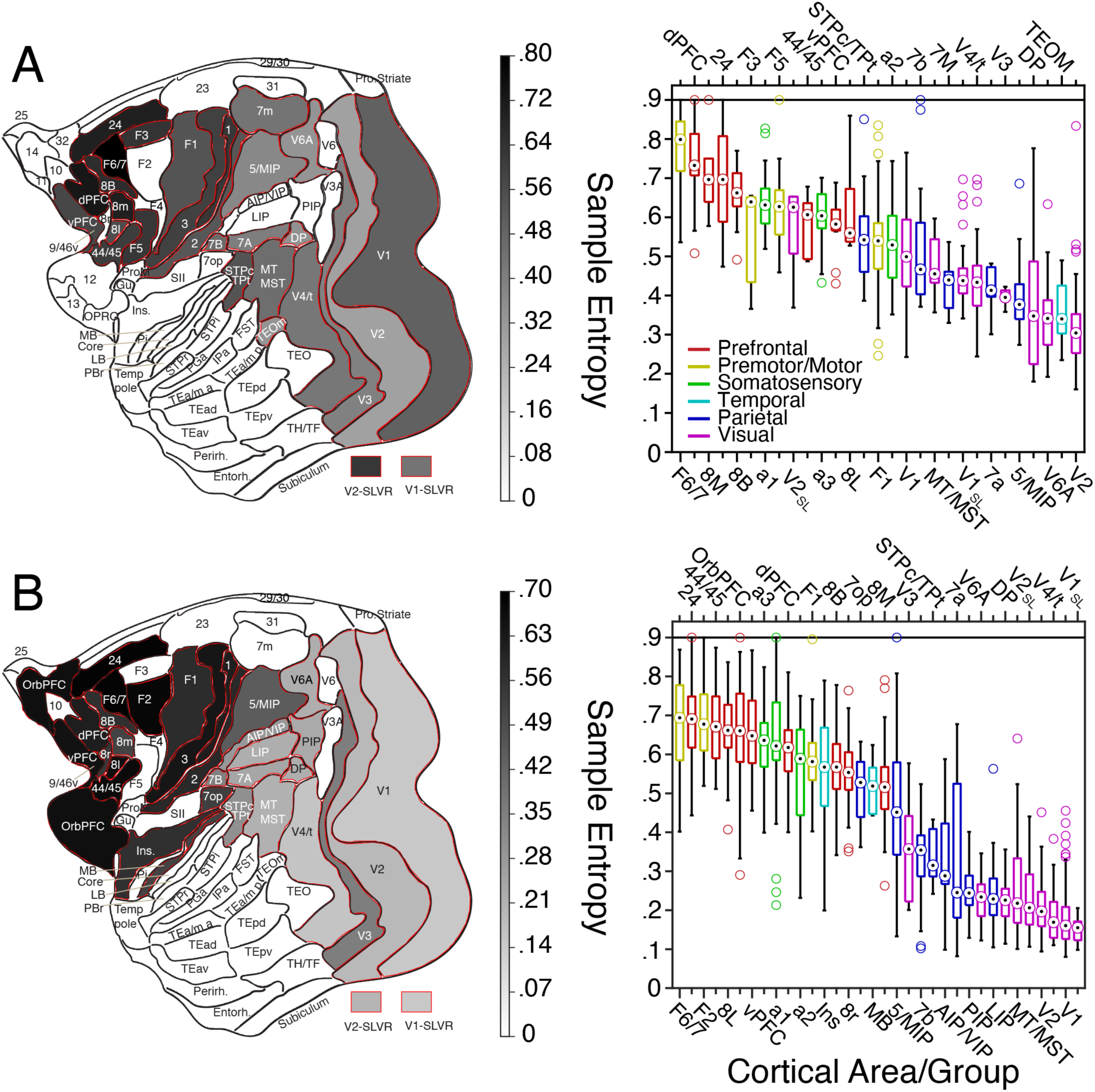
Sample entropy shows clear anatomical gradients. Variation of sample entropy (SampEn) across the cortex for both monkeys during the presample epoch. Cortical flatmaps of median SampEn (left) and corresponding distributions of SampEn, ranked by the median, for Monkeys E (A) and L (B). Plotting conventions are the same as in figure 3.

The organization of the LFP gradients suggested they may be related to the well-documented variation of synaptic spine counts on the basal dendrites of layer 3 pyramidal neurons (Elston, 2007), which exhibit a striking correlation to anatomical hierarchy (Markov et al., 2014b; Chaudhury et al., 2015; Wang, 2020). To test this conjecture, we computed the correlation between the published values of mean spine counts (Elston and Rosa, 1997, 1998a, b; Elston and Rockland, 2002; Elston et al., 1999, 2005, 2011) and the median values of SC, SampEn, and PA in each band in each monkey for all the cortical areas in which these data were available (Table 2). The distribution of average dendritic spine counts from a closely overlapping set of 14 areas in Monkey E and 18 areas in monkey L were correlated with the spectral content and peak amplitude in band-1 (SC1, PA1) and SampEn in both monkeys. Scatter plots and corresponding correlation coefficients (computed from all values in both monkeys) for these parameters are shown in figure 7. The low frequency spectral components (SC1, PA1) were negatively correlated with mean spine count while SampEn displayed a positive correlation. Significant correlations were also present in higher frequency bands in monkey L (SC2, SC3, SC5 and PA2) but not monkey E (Table 2).

**Figure 7.**
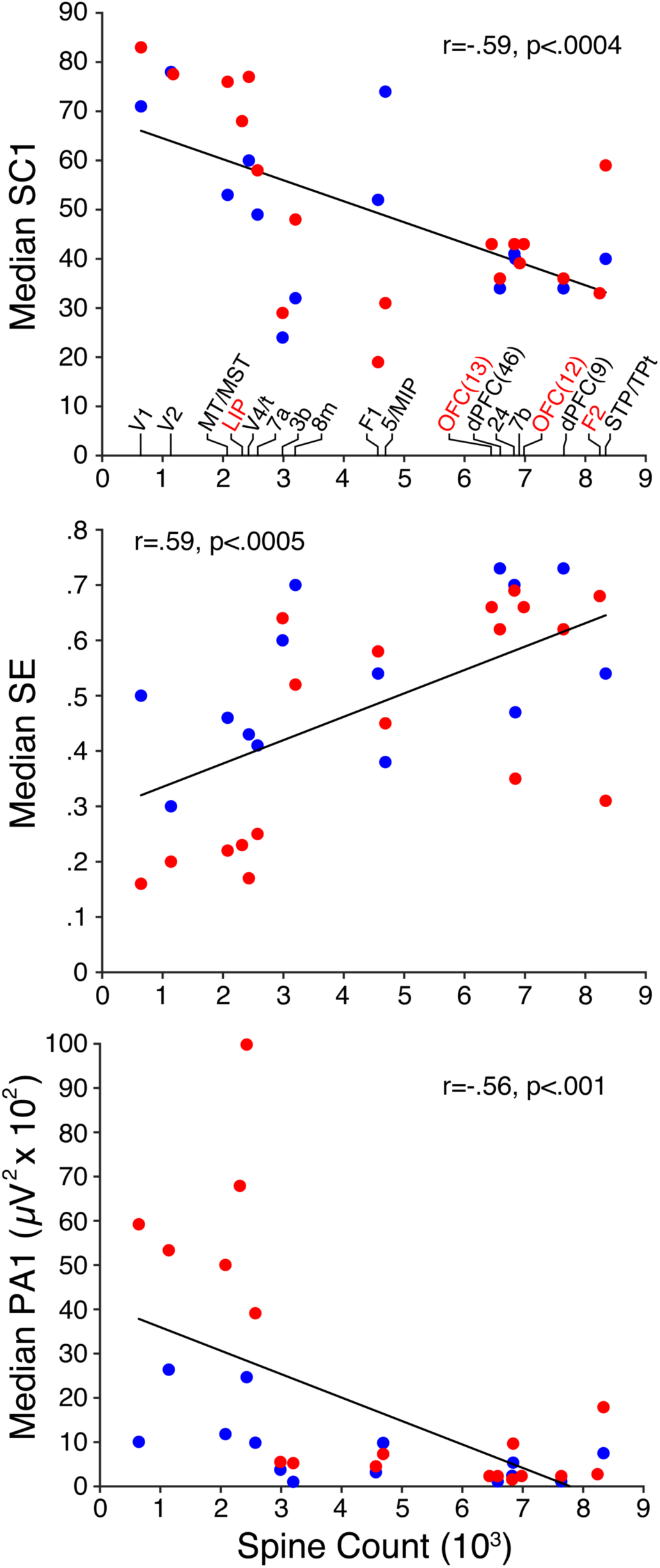
Scatter plots of the median values of SC1, SampEn and PA1 versus mean dendritic spine count on the basal dendrites of layer 3 pyramidal neurons for an overlapping set of 14 and 18 cortical areas in monkey E (blue) and monkey L (red). The correlation coefficients were calculated on the combined data from both monkeys. Spine counts along the x-axis are the same in all three plots and areal labels are shown above the x-axis in the top plot. Data for the areas labeled in black text were available for both monkeys while those labeled in red were available for monkey L only. Spine count data were taken from the reports of Elston and colleagues (Elston and Rosa, 1997, 1998a, b; Elston and Rockland, 2002; Elston et al., 1999, 2005, 2011). For some area groups (i.e., MT/MST, V4/t, 5/MIP, STP/TPt) spine data was obtained from just one area (i.e., MT, V4, 5, STP). For other area groups (i.e., OFC and dPFC) spine data was obtained from a subset of those areas (i.e., 12, 13, 9 and 46).

**Table 2.** Correlation coefficients, and corresponding P values, between the median value of each spectral parameter in each frequency band and mean spine count of the basal dendrites of layer 3 pyramidal neurons in an overlapping set of 14 areas in monkey E and 18 areas in monkey L (adopted from Elston and Rosa, 1997, 1998a, b; Elston and Rockland, 2002; Elston et al., 1999, 2005, 2011). Spectral parameters are derived from the presample epoch of the task. Numerical values 1-5 indicate the separate frequency bands for each monkey. Significant values are highlighted in red.

| **<u>Monkey E</u>** | <u>SC1</u> | <u>SC2</u> | <u>SC3</u> | <u>SC4</u> |  | <u>SampEn</u> |  | <u>PA1</u> | <u>PA2</u> | <u>PA3</u> | <u>PA4</u> |  |
| --- | --- | --- | --- | --- | --- | --- | --- | --- | --- | --- | --- | --- |
| Correlation | **-.55** | .36 | .44 | .51 |  | **.54** |  | **-.59** | -.27 | -.1 | -.06 |  |
| P value | **<.05** | <.21 | <.12 | <.06 |  | **<.05** |  | **<.03** | <.35 | <.74 | <.85 |  |
| **<u>Monkey L</u>** | <u>SC1</u> | <u>SC2</u> | <u>SC3</u> | <u>SC4</u> | <u>SC5</u> | <u>SampEn</u> |  | <u>PA1</u> | <u>PA2</u> | <u>PA3</u> | <u>PA4</u> | <u>PA5</u> |
| Correlation | **-.64** | **.6** | **.73** | .21 | **.51** | **.68** |  | **-.71** | **-.62** | -.39 | -.04 | -.1 |
| P value | **<.005** | **<.009** | **<.0007** | <.4 | **<.03** | **<.002** |  | **<.0009** | **<.006** | <.12 | <.89 | <.69 |

### Decoding Cortical Areas

Having identified a set of features that reveal spatial gradients of the spectral and temporal properties of the LFP, we sought to determine if these features could be used to classify cortical areas. We used a support vector machine (SVM) classifier algorithm, using a K-fold cross validation scheme (K = 5), and applied it separately to the full data set from each monkey, on each epoch of the task (see Methods). Data were included in the analysis if each area contained a minimum of 20 recordings (resulting in 11 areas for Monkey E and 28 areas for Monkey L; Table 1). The analysis included 9 features for Monkey E (SC and PA in 4 frequency bands, and SampEn) and 11 features for Monkey L (SC and PA in 5 frequency bands, and SampEn). The results of the classification analysis are shown in Figure 8. The confusion matrices (Figure 8A and B), and corresponding distributions of validation accuracies (VA) (Figure 8C and D), are shown for the presample epoch in monkey E (A and C) and monkey L (B and D). The median values ranged from 35%-85% in Monkey E (Figure 8A and C) and 20%-75% in Monkey L (Figure 8B and D) and all areas exceeded the 99% confidence limit, computed from the permutation test (red lines in Figures 8C and D, see Methods). As suggested by the gradients of SC, PA and SampEn, classification errors tended to lie near the diagonal, indicating similarity in the spectral and temporal properties of the LFP among nearby areas within a cortical region.

**Figure 8.**
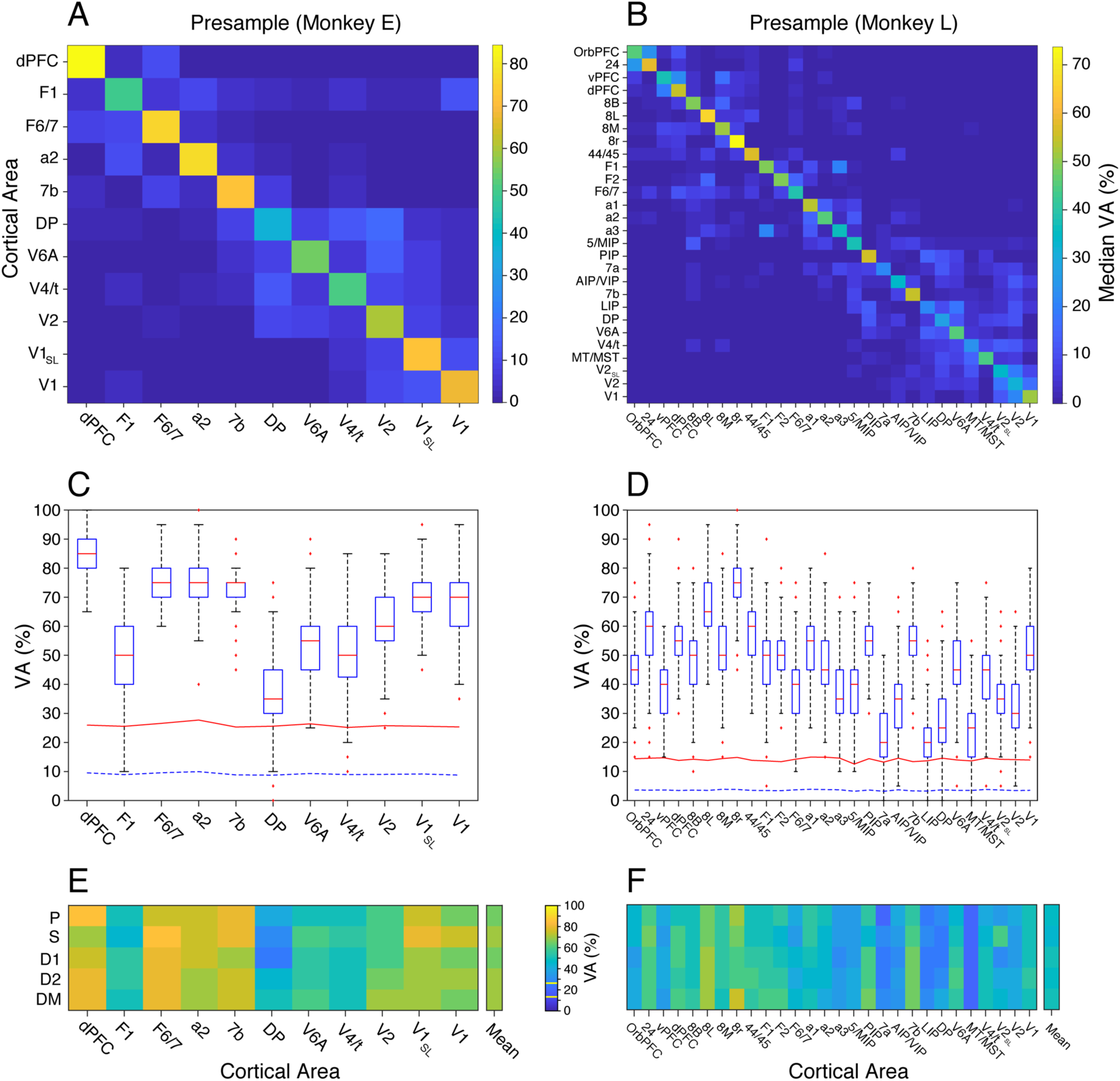
Results of the decoding analysis. Confusion matrices (A, B) and distributions of validation accuracies (C, D) for the presample epoch in monkey E (A, C) and monkey L (B, D). In each box plot the red line shows the median, the blue box displays the interquartile range, the whiskers show the 5^th^ and 95^th^ percentiles, and the red asterisks show outliers. The dashed blue line and solid red line in C and D show the mean and 99^th^ percentile computed from the surrogate distributions. The plots in E and F show the median validation accuracies as a function of task epoch (P, S, D1, D2, DM) for each area in monkey E and monkey L, respectively. The upper and lower yellow horizontal lines in the color-scale bar show the 99^th^ percentile computed from the surrogate distributions for monkey E and L, respectively.

To determine if VA was task-dependent, we compared the distributions in each area across the 5 epochs of the task. Figures 8E and 8F show the median VA in each area as a function of task epoch in monkeys E and L, respectively. These are the median values along the diagonals of the confusion matrices computed on each epoch. The corresponding distributions of VA are shown in figure 9. Task-dependent changes in VA were apparent in every area in both monkeys. In some areas VA increased during the delay period (e.g., DP and V2 in Monkey E; 7b and V6A in Monkey L), other areas displayed the opposite pattern (a2 in Monkey E; a1 and a2 in Monkey L), and many others displayed differences between epochs with no discernible pattern across areas. In fact, every area examined in each animal showed a significant difference in VA across epochs (Kruskal-Wallis test, P<<10^-5^, FDR corrected), suggesting widespread changes in the spectral and temporal properties of the LFP during the task that span all the cortical areas measured.

**Figure 9.**
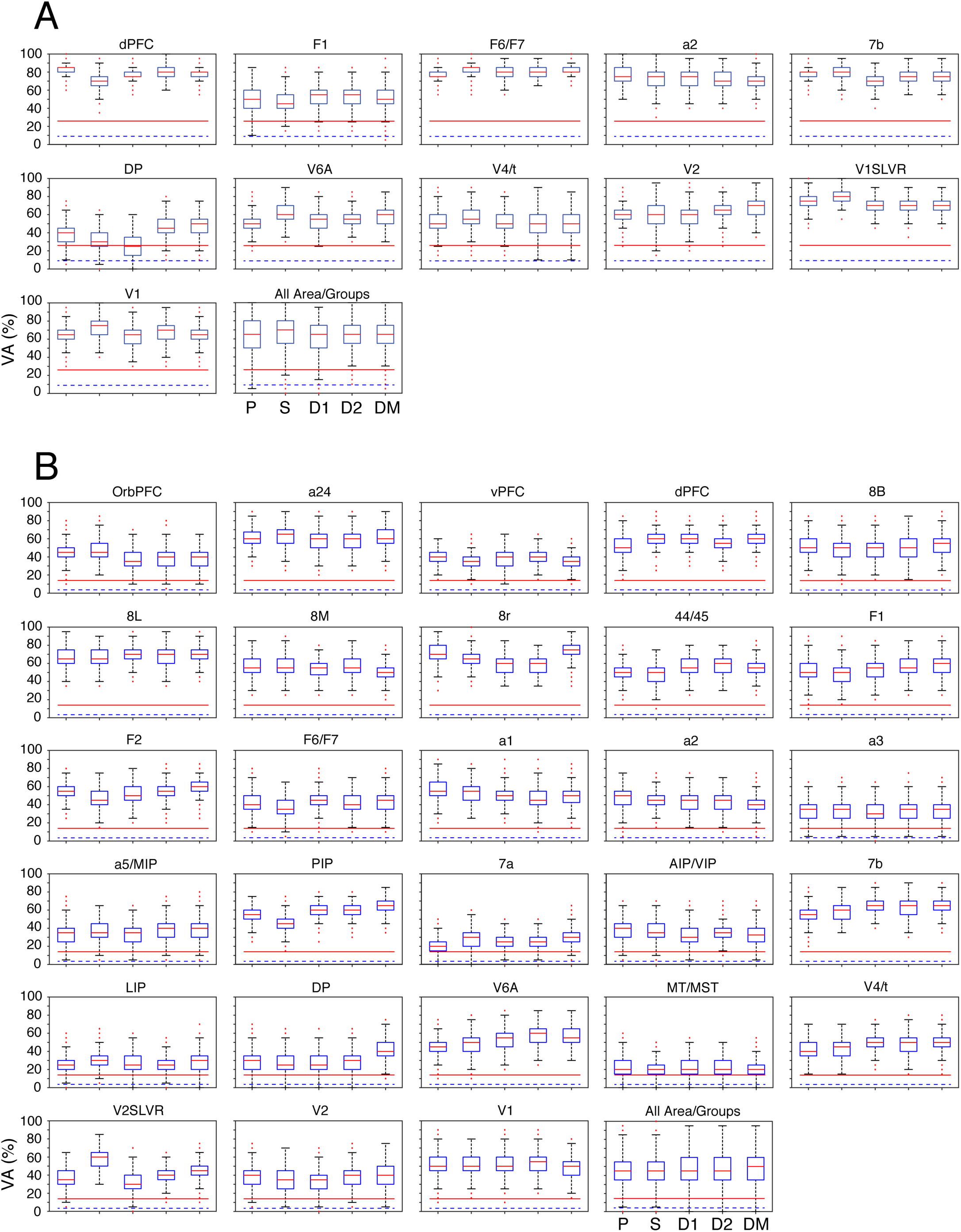
Distributions of validation accuracies (VA%) as a function of task epoch (P, S, D1, D2, DM) for each area in Monkey E (A) and Monkey L (B). The bottom right plot in A and B shows the cumulative distributions of validation accuracies across all areas. In each box plot the red line shows the median, the blue box displays the interquartile range, the whiskers show the 5^th^ and 95^th^ percentiles, and the red asterisks show outliers. The dashed blue line and solid red line in each plot show the mean and 99^th^ percentile computed from the surrogate distributions.

We performed several additional analyses to determine which features in the classifier were responsible for successful classification (see Feature Importance in Supplementary Materials). We first assessed the change in VA that occurs when the values of each feature are separately randomized (Dreyfus and Guyon, 2006). This led to a ∼5% decrease in the mean VA across areas for each feature in both monkeys as compared to the baseline (Figure S8). We ran two additional analyses, after excluding SampEn as a feature. We assessed VA independently for SC and PA using the corresponding features in all frequency bands and we assessed the effect on VA of removing both features in each frequency band separately (Tables S1 and S2). In both analyses, we found widespread, and often weak, effects that occurred in nearly all areas. We conclude that each feature makes a small contribution to validation accuracy and that no frequency band was substantially more informative than another. We also sought to determine which features of the spectra account for classification errors between areas (see Areal Misclassification in Supplementary Materials). We found an inverse relation between VA and the variance of SC and PA in low frequency bands (SC1, SC2, PA1, PA2 in Monkey E; SC1, SC2, PA1 in Monkey L) (Table S3) and a greater incidence of classification errors among nearby areas (Figure S9). Thus, both high variance in spectral features as well as similarity in feature values between nearby areas reduced validation accuracy. The latter result is consistent with our finding of spatial gradients in the spectral and temporal features of the LFP.

### Task Dependent Changes in Power

Although the classification analysis reveals widespread task dependent changes in the cortical LFP, it does not provide a direct measure of the incidence, magnitude, or sign of those changes for each cortical area. We therefore performed a separate analysis of the change in spectral power across task epochs and frequency bands for each recording in the data set used for the classification analysis (11 areas in monkey E, 28 areas in monkey L). To screen out signals with low amplitudes the mean spectral content in each band and epoch had to exceed 5%. For each trial in a session, we calculated the mean power within each band and each epoch. We compared the distribution of values across trials in the presample epoch to the distributions in each of the other four epochs using the Wilcoxon signed-rank test (p<.01, FDR corrected). This was repeated for every recording in an area resulting in a distribution of significant changes in power (both increases and decreases) occurring in each epoch and frequency band relative to the presample period (Figure 10). Example results are shown in figure 10A (area 8B (band-2) and area F2 (band-4) in monkey L). The mean of each distribution (black filled circle) and the incidence of significant differences (black horizontal line) are color coded and replotted in the lower two plots (see color scale in C and D). In these examples, there was a mixture of power increases and decreases in band-2 of area 8B that vary with task epoch, while the power in band-4 of area F2 shows a sustained and increasing suppression throughout the task epochs (note that the incidence of significant suppression in area F2 exceeds 90% in each epoch). The overall incidence of changes in power, across all areas, epochs, and frequency bands, is shown for both monkeys in figure 10B (left plot: monkey E; right plot: monkey L). Significant changes in LFP power, relative to the presample period, occurred in 60% and 50% of the epochs in monkeys E and L, respectively. Decreases in power occurred nearly twice as often as increases in both monkeys.

**Figure 10.**
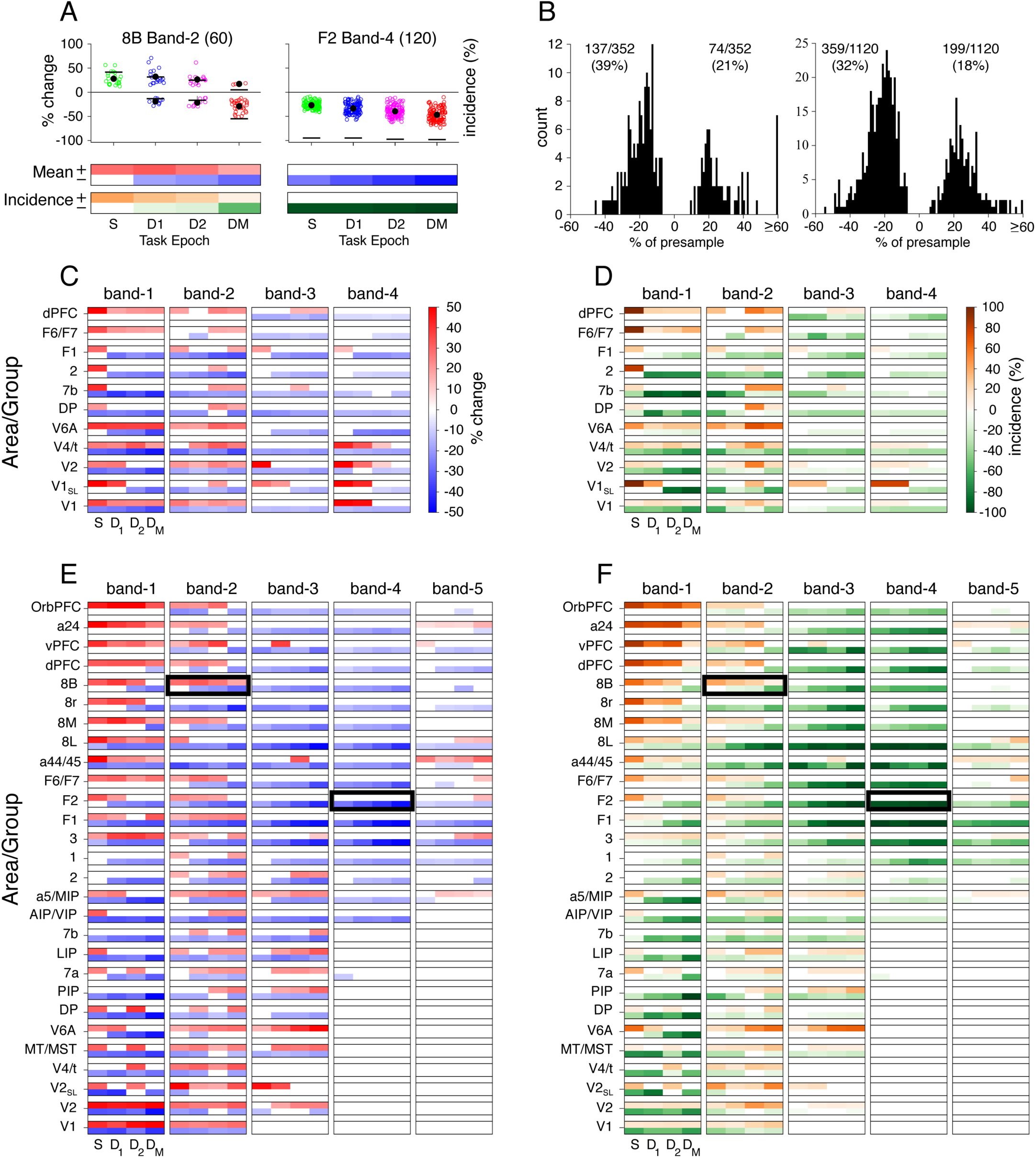
Task-dependent changes in LFP power as a function of task epoch, frequency band and cortical area in monkeys E and L. (A) Example results for area 8B (band-2) and area F2 (band-4) in monkey L. The upper plots show the distributions of significant changes in power across recording sites for each task epoch (S-sample, D1-delay1, D2-delay2, DM-delaym) as a percentage change relative to the presample epoch. The mean of each distribution is shown by the black filled circles. The incidence of significant values in each epoch is shown by the black horizontal lines. Recording counts are shown in parentheses. The mean change and the incidence values are color coded and displayed in the lower pair of plots for the two areas (mean change (blue/red), incidence (orange/green), see color scales in C and D). (B) Histograms of the change in mean power, relative to the presample epoch, for all task epochs, frequency bands and cortical areas in monkey E (left) and monkey L (right). The ratios in each plot show the incidence (%) of significant decreases and increases in power. (C-F) Summaries of the significant changes in mean LFP power (C, E) and the incidence of those changes (D, F) as a function of task epoch (S, D1, D2, DM), frequency band and cortical area for monkey E (C, D) and monkey L (E, F). The number of recordings in each area are given in Table 1. The black boxes in E and F indicate the data shown in the lower two pairs of plots in A.

To visualize the overall spatial pattern of these changes, we plotted the mean change in power and the incidence of occurrence in all epochs and frequency bands for all areas in monkey E (C, D) and monkey L (E, F). While the results differed between the two animals, several common findings were apparent in these plots. First, significant task-dependent increases and decreases in power occurred in at least one frequency band in every epoch of all areas in both monkeys, demonstrating widespread cortical involvement in the task. Second, the magnitude and incidence of the changes varied widely across areas and frequency bands. The spatial organization of the changes was most apparent in the incidence maps of Figure 10D and F. In band 1 prefrontal areas showed a task dependent enhancement, particularly in monkey L, while suppression was common and robust in central, parietal and occipital regions in both monkeys. In band 2 a complex combination of increases and decreases in power occurred across the cortical areas, making the spatial organization difficult to discern. Notably area V6A displayed a ramp-like enhancement in band 2 in both monkeys, while area 8L showed a progressive suppression in monkey L. In band 3 the most apparent pattern was a combination of suppression in somatomotor, premotor and prefrontal areas, and enhancement in parietal and extrastriate areas, particularly area V6A in monkey L. In band 4 of monkey L, which was unique to this animal (see Figure 3), a pronounced and ramp-like suppression of both the incidence and magnitude occurred throughout a broad region of cortex that included anterior parietal, somatomotor, premotor, and prefrontal areas. At the higher frequencies (band 4 in monkey E, band 5 in monkey L), the pattern of task-dependent changes differed between the two animals. In monkey E there was a distinct enhancement in response to the sample stimuli in early visual cortex, most notably in foveal and perifoveal V1 (V1-SLVR). In monkey L the changes in band 5 showed a close spatial correspondence to the changes in band 4, but the responses were a mixture of increases and decreases of activity. Together these results demonstrate widespread, task-dependent changes in the cortical LFP that span all the areas we recorded from. Moreover, the spatial organization of these changes varies with frequency and task epoch.

## DISCUSSION

We made simultaneous measurements of intracortical neural activity in two monkeys performing a visual short-term memory task (Dotson et al., 2017, 2018) to investigate the spatial organization and task-dependence of the LFP across a significant fraction of the anatomically identified cortical areas in non-human primates (Markov et al., 2014a). Analysis of the peak frequencies in the LFP power spectra revealed multiple narrow frequency bands in both animals. These distributions overlapped, but also differed in some key respects, demonstrating that the spectral composition of the cortical LFP can vary significantly between subjects and does not always fall neatly into classically defined canonical frequency bands. These results, along with differences in the recording locations and sample sizes, required us to confine the subsequent analyses to each monkey individually. We found that spatial maps and rank-ordered plots of two spectral parameters (SC, PA) revealed spatial gradients and large amplitude differences across the cortical map that differed markedly between frequency bands. These gradients were similar in the two monkeys, apart from band-4 in monkey L that was absent in monkey E. Separate analysis of the temporal complexity of the LFP, using the measure of Sample Entropy (SampEn), revealed a similarly striking frontal to occipital gradient across the cortical map in both monkeys (Figure 6). Together these analyses demonstrate clear areal differences in the spectral and temporal properties of the LFP that exhibit multiple, distinct spatial gradients across the cortical map. These gradients are reminiscent of the earliest qualitative reports in humans and monkeys (Jasper and Andrews 1938; Garvin and Amador 1949; Jasper and Penfield 1949) and exhibit some similarities to a recent report in humans (Frauscher et al., 2018).

We also compared the spectral and temporal metrics of the LFP to the well documented gradients of synaptic spine counts on the basal dendrites of layer 3 pyramidal neurons (Elston and Rosa, 1997, 1998a, b; Elston and Rockland, 2002; Elston et al., 1999, 2005, 2011), which are closely correlated with areal hierarchy defined by interareal connections (Markov et al., 2014a,b; Chaudhury et al., 2015). This analysis revealed significant correlations between the synaptic spine counts and the low frequency spectral components (SC1, PA1) and SampEn (Figure 7) in both monkeys. Significant correlations were also present for SC and PA in bands 2 and 3 in monkey L. These findings are particularly interesting, given the central role of temporal summation of dendritic synaptic currents in the genesis of the cortical LFP (Linden et al., 2011; Einevoll et al., 2013; Pesaran et al., 2018). However, we do not understand how the patterns of synaptic current give rise to the different spectral components of the LFP, nor how variations in synaptic organization might contribute to the gradients we observe. These findings are a new piece in a larger puzzle requiring further research.

Using the measures of spectral power and SampEn, we could reliably classify cortical areas, or small groups of areas, well above the 99% confidence limit derived from surrogate distributions that randomized areal assignment. The validation accuracy varied across epochs of the task in all areas. However, no single feature or frequency band incorporated in the analysis stood out as particularly informative, and validation accuracies exhibited a wide range in both monkeys. This suggested that variance in the feature distributions, or similarities in those distributions among nearby areas, could have led to degraded classification performance. We found evidence supporting both of these conjectures. Together these analyses demonstrate widespread task-dependent changes in the LFP and show that the spectral features of the LFP display a considerable degree of overlap among adjacent areas, indicating that the LFP varies at a regional level incorporating multiple areas with related functional properties (Keitel and Gross, 2016).

The decoding analysis, however, provided little or no information regarding the specific changes in spectral power that occur in each area during the task. We therefore calculated the magnitude and incidence of changes in power that occur in each area with respect to each frequency band and epoch of the task. This revealed the striking result that every area of the cortex we sampled in both monkeys displayed both significant increases and decreases in LFP power in multiple frequency bands and multiple epochs of the task. Thus, even a simple cognitive task, such as remembering a salient visual object for 1-2 seconds, evokes changes in power across widespread areas of the neocortex spanning, visual, parietal, somatomotor, premotor and prefrontal areas. This result is consistent with our earlier findings of widespread, task-dependent changes in unit activity in the same data set (Dotson et al., 2018). These findings are also consistent with a functional imaging study in humans demonstrating task-dependent changes in the BOLD signal in widespread regions of the brain during a simple visual attention task (Gonzalez-Castillo et al., 2012).

### Methodological Considerations

Our approach represents a significant advance over other methods such as ECoG that are limited to surface measurements with lower spatial resolution (Bosman et al., 2012; Bastos et al., 2015; Dubey and Ray, 2019). However, the study also had several methodological limitations. First, we were unable to record from most ventral areas of the cortex, particularly in the temporal lobe. Second, the cortical areas we recorded from, and the size of the sample measured from each area, differed substantially between monkeys and between areas within each monkey. Part of this was due to an improvement in the methods used in Monkey L (Dotson et al., 2017). This sampling problem could have biased our results. For example, we obtained a large sample of recordings in area F2 in Monkey L, but none in Monkey E (Table 1). We made efforts to mitigate this problem by analyzing the data from each monkey separately and by sub-sampling the data from each area in each iteration of the decoding analysis. Third, although we were able to recover the areal location of each recording, we could not identify the cortical layer at each recording site. Given that the LFP is known to vary in amplitude and frequency across the cortical layers (Murthy and Fetz, 1996; Spaak et al., 2012; Bastos et al., 2018), the lack of layer information could have introduced significant variance or biased our results. Fourth, variations in arousal level are known to have pronounced influences on the spectral properties of the LFP (Magoun, 1961), and these were clearly present in our data, particularly when the animals became drowsy or disinterested in the task (see Figure 11). We attempted to reduce these effects by restricting our analysis to correct trials during periods of high performance on the task. Finally, the type of cognitive task, and the stimuli used in the task, are likely to have a significant influence on the properties of the LFP that depend on cortical area.

**Figure 11.**
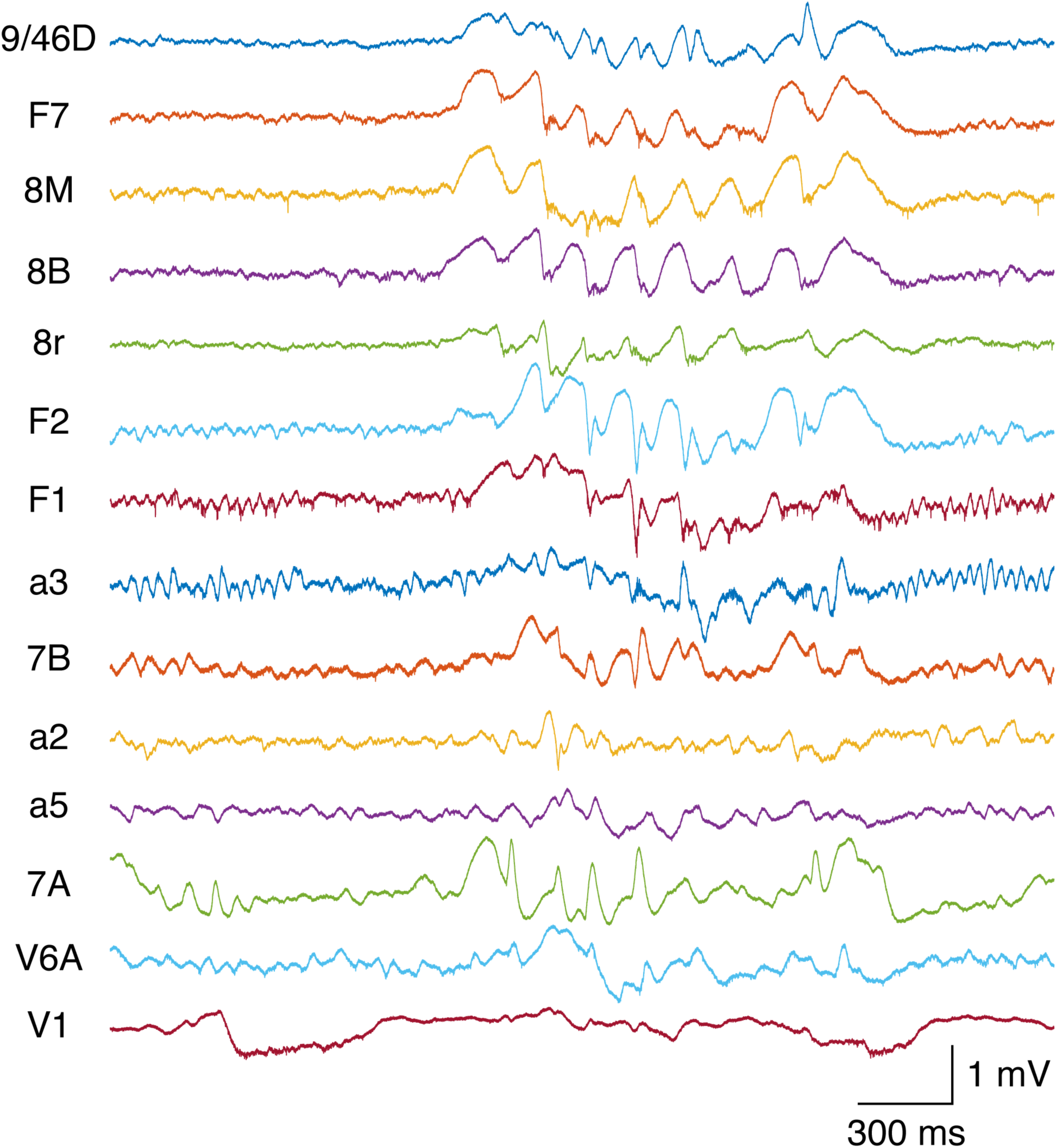
Plots of a 3-second segment of broadband raw data recorded during a period of rest with the room lights turned off in monkey L. The data is shown for the same channels on the same session as the plots in figure 1. A rapid-onset sleep spindle occurs halfway into the segment with high amplitude in prefrontal, premotor, and parietal areas.

There were three other factors that could have affected our ability to discriminate differences in the LFP between cortical areas. First, because we used monopolar recording methods, volume conducted signals could have blurred the differences between areas. However, multiple lines of evidence indicate that the LFP is highly local (Gray and Singer, 1989; Katzner et al., 2009; Spaak et al., 2012; Einevoll et al., 2013; Pesaran et al., 2018; Dubey and Ray, 2019), suggesting that volume conduction has minimal effects on our data. Second, the reference channel in our recordings was tied to the large titanium chamber implanted on the animals (Dotson et al., 2017). This provided a “quiet” reference potential by integrating signals from widespread areas of the skull. A separate analysis of the spectral coherence between simultaneously recorded LFPs revealed many instances of non-significant coherence at all frequencies (data not shown), suggesting that the reference signal is indeed quiet. Third, the 400 ms duration of the epoch-based spectral analysis, the multi-taper filtering of the power spectra, and the AC-coupling of our recording system, limited our ability to evaluate signals less than 4 Hz.

Finally, a further analytical issue concerns the spectral and temporal metrics we chose for characterizing the LFP. The SC and PA metrics provide overlapping and somewhat redundant information. However, they also made separate and useful contributions to the decoding analysis. In nearly all cortical areas, one feature performed significantly better than the other (see Tables S1 and S2). This can occur, for example, when there is a big difference between areas in absolute power (PA), while the differences in relative power (SC) remain small. A related issue is present with SampEn, where the value of the metric likely reflects the relative power of low and high frequencies. In spite of this apparent redundancy, SampEn was informative in classifying cortical areas (see Figure S8). This metric also revealed a clear occipito-frontal gradient in both animals despite significant differences between animals in the distribution of peak frequencies shown in figure 3 and notable differences in the distribution of peak amplitudes shown in figure 5. Such a result would not be expected if this metric only reflected the relative magnitudes of low and high frequency power.

### Frequency Differences

Outside of the low frequencies in band-1, we found several notable differences in the distribution of spectral peaks in the two monkeys. Signals with spectral peaks in band-2 were widely distributed across the cortical areas in both monkeys but the frequencies were centered at 14 Hz in monkey E and 10 Hz in monkey L. Signals with spectral peaks in band-3 were were frontally distributed in both animals and centered at ∼25 Hz in Monkey E and ∼20 Hz in Monkey L. The narrow range of spectral peaks in bands 4 and 5 of Monkey L appeared to be completely absent in Monkey E. Part of these differences may stem from the sampling problem discussed above. But perhaps the simplest explanation is that the occurrence and frequency distribution of narrow band oscillatory activity can differ widely between individuals and do not necessarily fall into the classically defined canonical frequency bands. This conclusion is supported by studies in both monkeys and humans (Kilavik et al., 2012; van Pelt et al., 2012; Confais et al., 2020; Vezoli et al., 2021), where the frequencies of narrow band oscillatory activity can differ significantly between individuals.

Regarding the higher frequency bands (band-4 in Monkey E and bands 4 and 5 of Monkey L), several results from this study were particularly noteworthy. First, the spectral peaks in band-5 of Monkey L (Figures 2 and 3) co-occurred at twice the frequency of the high amplitude narrow-band oscillations in band 4, suggesting a higher harmonic that is unlikely to reflect a distinct spectral component. Our supplemental analysis (Figures S4-S7) confirmed this conjecture. Second, visually evoked gamma-band (30-80 Hz) oscillations, characteristic of primary and early extrastriate areas of the visual cortex (Gray and Singer, 1989; Friedman-Hill et al., 2000; Fries et al., 2001; Bosman et al., 2012; Bartoli et al., 2019), were notably sparse and low amplitude in our data. This type of activity is known to depend on the use of appropriate visual stimuli that are tailored to the receptive field properties of the cortical neurons in those areas. While we did observe clear instances of visually evoked gamma-band oscillations in V1 and V2, particularly in Monkey E, this was largely by chance, since we made no attempt to measure cellular receptive fields and adjust the sample stimuli to optimally activate the neurons at those recording sites. Third, we found limited evidence of elevated activity at the higher frequencies in either monkey during the delay period of the task (Figure 10). In contrast, the oscillatory activity in this frequency range was nearly universally suppressed during the memory delay. This finding differs from several studies reporting elevated gamma-band activity during the delay period of short-term memory tasks (Pesaran et al., 2002; Howard et al., 2003; Meltzer et al., 2008; Roux et al., 2012; Honkanen et al., 2015; Lundqvist et al., 2016), and argues against a role of gamma-band activity as a general information carrier in the neocortex (Fries, 2009; Basar, 2013).

### Implications for Cortical Processing

Our findings also raise important questions regarding the concept of a hierarchy of intrinsic timescales across the cortex. The prevailing view, based on functional imaging and ECoG studies in humans (Hasson et al., 2008, 2016; Honey et al., 2012; Gao et al., 2020) and single unit studies in monkeys (Murray et al., 2014; Dotson et al., 2018; Cavanagh et al., 2020; Spitmann et al., 2020), indicates that the intrinsic timescale, and temporal receptive window (Hasson et al., 2015), of cortical areas gradually increase across the cortical hierarchy from sensory to association areas. Correspondence between the functional imaging and single unit measures has also been recently established in monkeys (Manea et al., 2022). Our findings, however, suggest an additional perspective. We find multiple, overlapping gradients of LFP power across cortical areas that display striking differences with respect to frequency (Figures 4 and 5). Occipital-to-prefrontal gradients at low frequencies (band 1) transition to prefrontal-to-occipital gradients at higher frequencies (bands 3-5), while intermediate frequencies (band 2) display a unique gradient radiating from parietal and sensory-motor areas. If we consider 1/frequency as a simple measure of time scale, these gradients differ from and may be superimposed upon the hierarchical gradients established in previous studies. These findings may also account for the diversity of intrinsic time scales described in recent single unit studies (Cavanagh et al., 2020; Spitmann et al., 2020). Our finding of widespread task dependence of LFP power further suggests that these gradients dynamically change with respect to cognitive task, attentional demands and the types of sensory stimuli and motor actions (Gao et al., 2020).

Finally, our findings have important implications for the enduring view of cortical organization and function proposed by Mountcastle (1979). In that classical monograph, Mountcastle proposed two fundamental tenets of cortical organization: The first of these, the unit module, synonymous with the cortical column, was proposed as a canonical circuit in which “*… the processing function of neocortical modules is qualitatively similar in all neocortical regions … without the appearance of qualitatively different modes of intrinsic organization*”. The striking differences we and many others find in the spectral and temporal properties of the LFP across cortical areas do in fact reveal “*qualitatively distinct modes of intrinsic dynamics*”. When these results are combined with the well documented anatomical and functional gradients across the cortex (Elston, 2007, 2011; Collins et al., 2010; Hasson et al., 2008, 2015; Honey et al., 2012; Murray et al., 2014; Goulas et al., 2018; Huntenburg et al., 2018; Hilgetag et al., 2019; Wang, 2020; Li and Wang, 2022; Froudist-Walsh et al., 2023), a strictly canonical columnar model of neocortex (Hawkins, 2021) appears untenable. The second tenet was the concept of the distributed system, whereby “… *complex function controlled or executed by the system is not localized to any one of its parts. The function is a property of the dynamic activity within the system: it resides in the system as such.”* Our finding of widespread task-dependent changes in spectral power that occur even for a simple cognitive task (Gonzalez-Castillo et al., 2012) reinforces the view of dynamic distributed processing as an integral feature of the central nervous system (Tognoli and Kelso, 2014).

## Supplemental Data

See supplementary materials online.

## Acknowledgments

We are grateful to Dr. Chris O’Rourke for her excellent veterinary care of the animals and assistance in this study. We thank Susan Krueger for her help in the behavioral training and daily care of the animals. We are grateful to Drs. Martin Vinck, Joachim Gross and Pedro Maldonado for their helpful comments on earlier versions of the manuscript. We also thank Neuralynx (Neuralynx, Inc., Bozeman MT, USA) for providing the data acquisition system.

## Author Contributions

S.J.H., N.M.D. and C.M.G. conceived and designed research, performed the surgical procedures, collected and analyzed the data, and wrote the manuscript. V.L. and C.M.G. performed the gamma frequency harmonic analysis.

