## Supplemental Data for "The Primate Cortical LFP Exhibits Multiple Spectral and Temporal Gradients and Widespread Task-Dependence During Visual Short-Term Memory"

### Supplementary Materials

#### Spectral Peak Detection

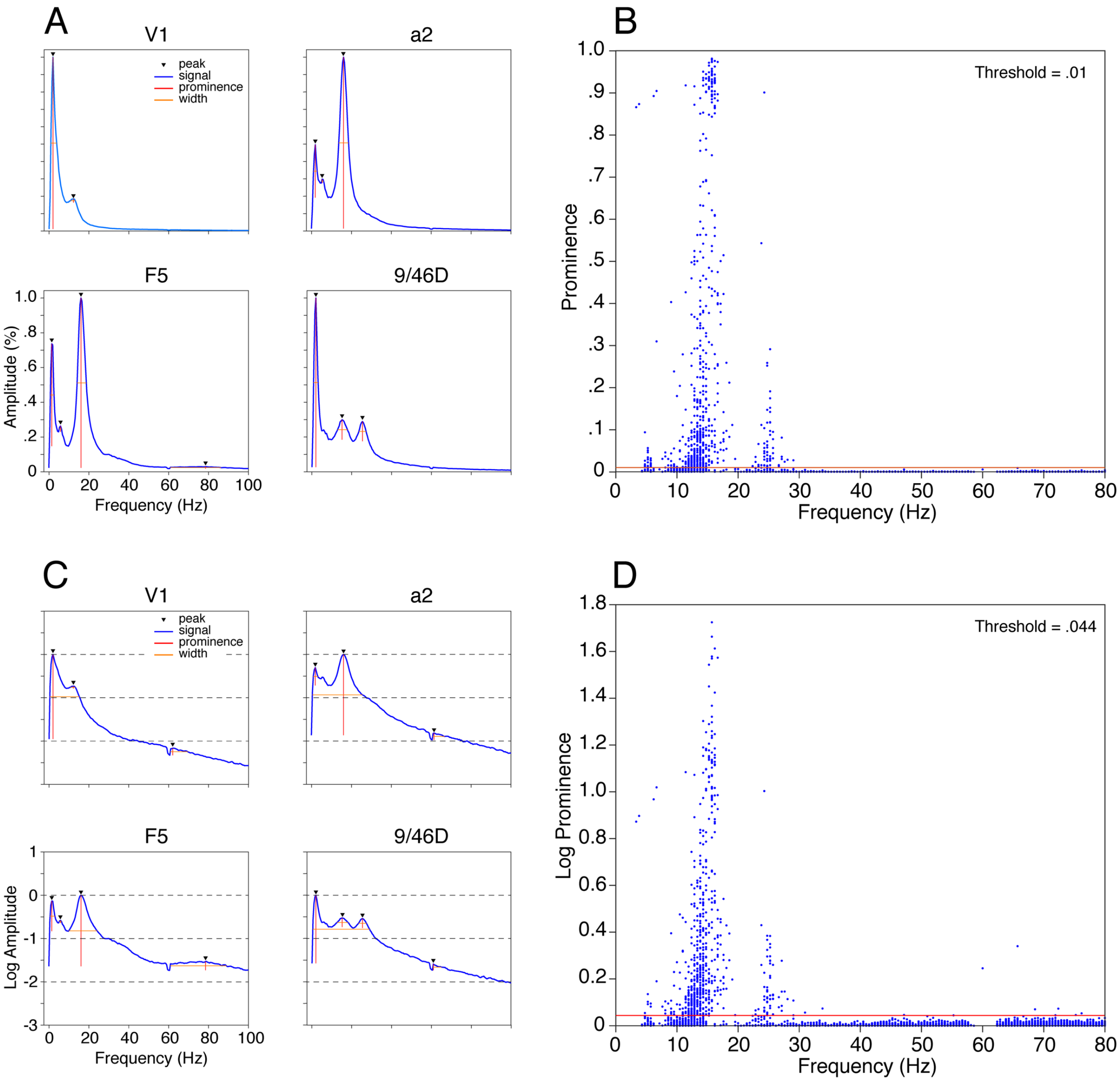

**Figure S1.** Peak detection results in Monkey E. (A, C) Examples of normalized average power spectra computed on the full trial duration (2.1s, see Methods) using all correct trials in a single session and displayed in linear (A) and semi-log (C) coordinates. Detected peaks are indicated by a filled triangle and the prominence and width of each peak are marked by red vertical and horizontal lines, respectively. Labels at the top of each plot indicate the cortical area from which the data were obtained. (B, D) Distributions of peak prominence as a function of frequency for all channels in all sessions obtained from the power spectra in linear (B) and semi-log (D)

coordinates, respectively. Peaks <4Hz were excluded from the analysis. The red horizontal line in each plot shows the peak prominence threshold used to exclude small peaks at the noise level.

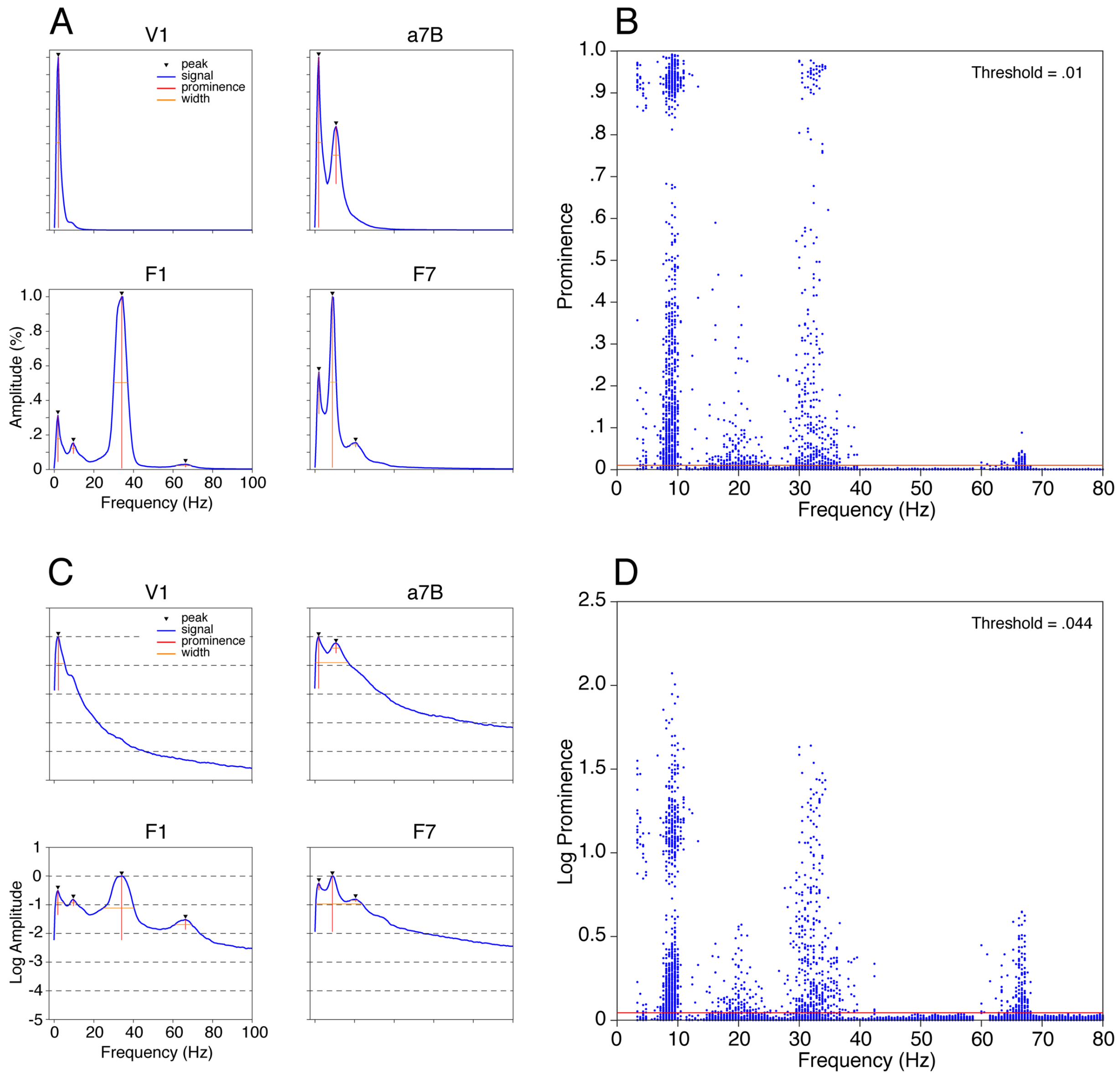

**Figure S2.** Peak detection results in Monkey L. (A, C) Examples of normalized average power spectra computed on the full trial duration (2.1s, see Methods) using all correct trials in a single session and displayed in linear (A) and semi-log (C) coordinates. Detected peaks are indicated by a filled triangle and the prominence and width of each peak are marked by red vertical and horizontal lines, respectively. Labels at the top of each plot indicate the cortical area from which the data were obtained. (B, D) Distributions of peak prominence as a function of frequency for all channels in all sessions obtained from the power spectra in linear (B) and semi-log (D) coordinates, respectively. Peaks <4Hz were excluded from the analysis. The red horizontal line in each plot shows the peak prominence threshold used to exclude small peaks at the noise level. Peak detection results in Monkey L. Descriptions and conventions are the same as those in figure S1.

**Distribution of Spectral Peaks**

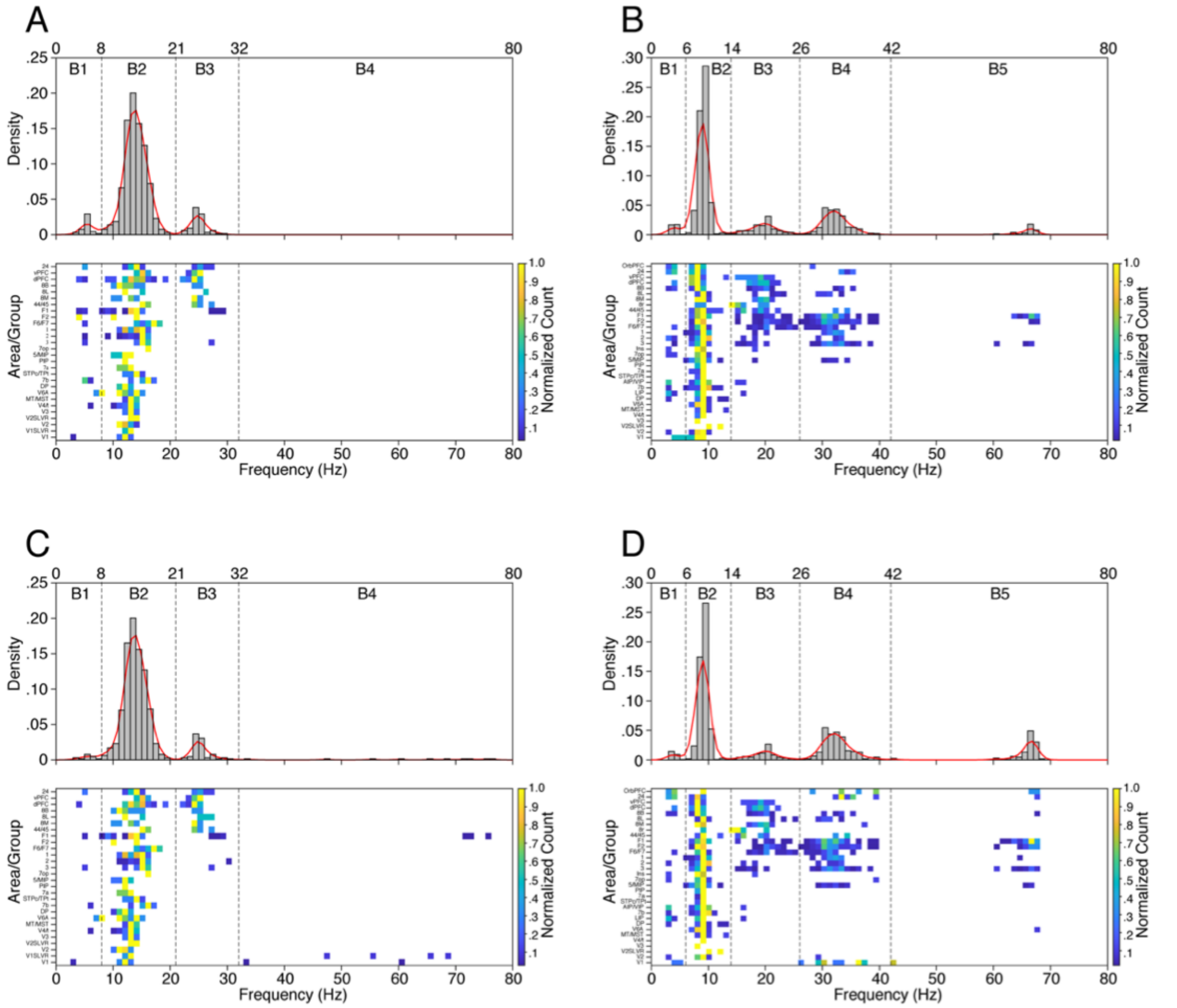

**Figure S3.** Distributions of spectral peaks obtained from all channels and sessions in Monkey E (A, C) and Monkey L (B, D) in linear (A, B) and semi-log (C, D) coordinates, respectively. The descriptions and conventions are the same as those in figure 3. The data in C and D are duplicated from figure 3 to facilitate comparison. There were small differences in the distribution of detected peaks derived from the linear and semi-log spectra, but these had no effect on the selection of frequency bands.

**Frequency Harmonic Analysis**

Two findings from the peak frequency histogram in monkey L (Figure 3B, Figure S3B,D) suggest that the band-5 oscillations reflect a higher order harmonic of the oscillations in band-4, and not an independent spectral component. First, with one exception, the band-5 spectral peaks occurred only in areas with a prominent peak in band-4. Second, the distribution of peak frequencies in band-5 was centered at approximately twice that of the band-4 peak frequency distribution. Higher order harmonics are known to occur in EEG, MEG and LFP data and are thought to result from the non-sinusoidal nature of the dominant oscillatory activity (Aru et al., 2015; Lozano-Soldevilla et al., 2016; Gerber et al., 2016; Cole et al., 2017; Schaworonkow and Nikulin, 2019). To further test this hypothesis, we sought to obtain answers to the following questions:

1) Are the temporal envelopes of the band-4 and band-5 spectral components correlated on each trial? If so, does the magnitude of the correlation depend on the amplitude of the band-4 oscillations?
2) What is the instantaneous phase difference between the band-4 and band-5 temporal envelopes and how does it vary with the instantaneous joint amplitude of the signals?
3) Are the band-4 oscillations non-sinusoidal in shape? If so, does this property change with the amplitude of the band-4 activity?
4) Are the band-5 oscillations phase locked to the band-4 oscillations?

To address these questions, we performed the following analyses on all the LFP signals in monkey-L that contained peaks in both band-4 and band-5 of the average spectrum that exceeded the peak prominence criterion.

To assess the temporal correlation and phase relationship between the band-4 and band-5 spectral components, we first computed the time-frequency (TF) spectrum of the LFP recorded on each trial (2.15 sec duration, ranging from -0.65s to 1.5s relative to sample onset) using the multi-taper method with a time-bandwidth product of 4 (Gramfort et al., 2013; Babidi and Brown, 2014). On each trial, we extracted the temporal envelope of the band-4 and band-5 signals by taking the moving average of the TF spectral values between 28-43 Hz and 58-74 Hz, respectively (temporal resolution = 10 ms). We refer to the resulting signals as b4(t) and b5(t). We then calculated the Pearson's correlation coefficient (CC) between b4(t) and b5(t) on each trial. Example results for a single trial are shown in Figure S4, which reveals a close temporal correlation. In this session, 7 channels displayed peaks in band-4 and band-5 of the average spectrum. The CC distributions for these channels are shown in Figure S5A. The observed distributions were greater than the corresponding trial-shuffled surrogate distributions in all 7 channels (KS-test, $p<<.001$). We next plotted the median CC versus the mean power in band-4 for all selected channels in the full data set (Fig. S5B) (n=132). This revealed that the strength of the correlation between b4(t) and b5(t) was proportional to the amplitude of the band-4 oscillations. The observed CC distribution exceeded the corresponding surrogate distribution on 129 out of 132 recordings (KS-test, $p<.001$).

To assess how the phase difference between b4(t) and b5(t) varies with the combined amplitude of the signals, we first normalized each signal to its respective z-score on each trial. We then computed the instantaneous correlation strength between the signals, $CC_{4,5}(t)$, using the method described in Faskowitz et al. (2020),

$$CC_{4,5}(t) = \frac{Z_4(t)Z_5(t)}{N-1}$$

where

N = number of samples on each trial

and the instantaneous phase difference on each trial, $\Delta\Phi_{4,5}(t)$, by taking the arctangent of the complex values of the Hilbert transform (Tass, 1998):

$$\Delta\Phi_{4,5}(t) = \Phi_4(t) - \Phi_5(t)$$

Using the results of these calculations, we plotted $CC_{4,5}(t) vs \Delta\Phi_{4,5}(t)$ for all trials and times on the selected channels in the data set (n=132). This revealed that the distribution of $\Delta\Phi_{4,5}(t)$ decreased as the magnitude of $CC_{4,5}(t)$ increased (Figure S5C). This result is further summarized by plotting the standard deviation of $\Delta\Phi_{4,5}(t)$ for discrete intervals of $CC_{4,5}(t)$ (Figure S5D). The variance of the phase differences decreases sharply as the magnitude of the correlation increases. Thus, b4(t) and b5(t) are closely correlated and the magnitude of the correlation is closely coupled to the joint amplitude of the signals.

Next, we sought to determine if the band-5 oscillations arise from the non-sinusoidal shape of the band-4 oscillatory activity. To test this hypothesis, we filtered the LFP signal in three different passbands: broadband (15-140 Hz) which removes the low frequency components, lowpass (15-50 Hz) which isolates the band-4 component, and highpass (50-80 Hz) which isolates the band-5 component. Examples of the resulting signals are shown Figure S6, which is taken from the same trial shown in Figure S4. The upper and middle plots show the filtered raw data in each of the three passbands at low and high temporal resolutions, respectively. The broadband signal (blue) has a non-sinusoidal waveform that is not readily apparent in the lowpass signal (black). Moreover, the highpass signal (red) displays a consistent phase relationship to both the broadband and lowpass signals.

To further examine these effects, we computed the cycle-triggered average (CTA) of these signals, using the time of the negative peaks in the broadband signal as the trigger points for averaging (indicated by the black filled circles in the middle plot of Figure S6). The LFP data were z-score normalized on each channel using the session mean and standard deviation before computing the CTA. This enabled us to easily compare the waveforms of the CTAs computed for different amplitude ranges of the LFP. The CTAs shown in Figure S6B were computed from the negative peaks ranging from 2-3 standard deviations in amplitude. The broadband CTA waveform (blue) appears less sinusoidal than the lowpass CTA (black). The highpass CTA (red) is tightly phase-locked to the negative peaks in the broadband signal. We measured the time lag of the highpass CTA on all recordings (n=132) and a histogram of the distribution is shown in Figure S6C. This reveals a close and consistent phase locking of the highpass signal to the negative peaks of the broadband signal across all recordings containing spectral peaks in both band-4 and band-5.

Finally, in order to assess the non-sinusoidal shape of the band-4 oscillations and its relation to the amplitude of those signals, we applied a simple metric of waveform asymmetry (ΔCT) to the CTAs computed from the broadband signals (Schaworonkow and Nikulin, 2019).

$$\Delta CT = \frac{T_C - T_T}{T_C + T_T}$$

where

$T_C$ and $T_T$ are the durations of the crests and trough periods in the CTA waveform, respectively (Fig. S7A). When the values of $T_C$ computed on either side of the trough differed, we chose the average of the two values for calculating ΔCT. We computed ΔCT for 3 different amplitude ranges of the highpass signal (1-2 std, 2-3 std, 3-4 std) and compared the resulting values to the amplitude of the band-4 power.

The results of these calculations, shown in Figure S7, reveal several findings. On average, waveform asymmetry increases with the mean power in band-4, and the slope of this relationship scales with the amplitude of the waveforms used to compute the CTA. These effects are shown for the example case in Figure S7A, and the general trend is apparent in Figure S7C. However, the effect is mixed and does not hold for those channels in which band-4 power is on the low end of the scale. This is illustrated by the example in Figure S7B and for those values to the left of the vertical dashed lines (<1) in Figure S7C. Replotting the data after excluding the channels with low power in band-4 reveals the trend (Fig. S7D).

In summary, our calculations demonstrate that band-4 oscillatory activity exhibits non-sinusoidal waveforms that give rise to a higher harmonic spectral component in band-5.

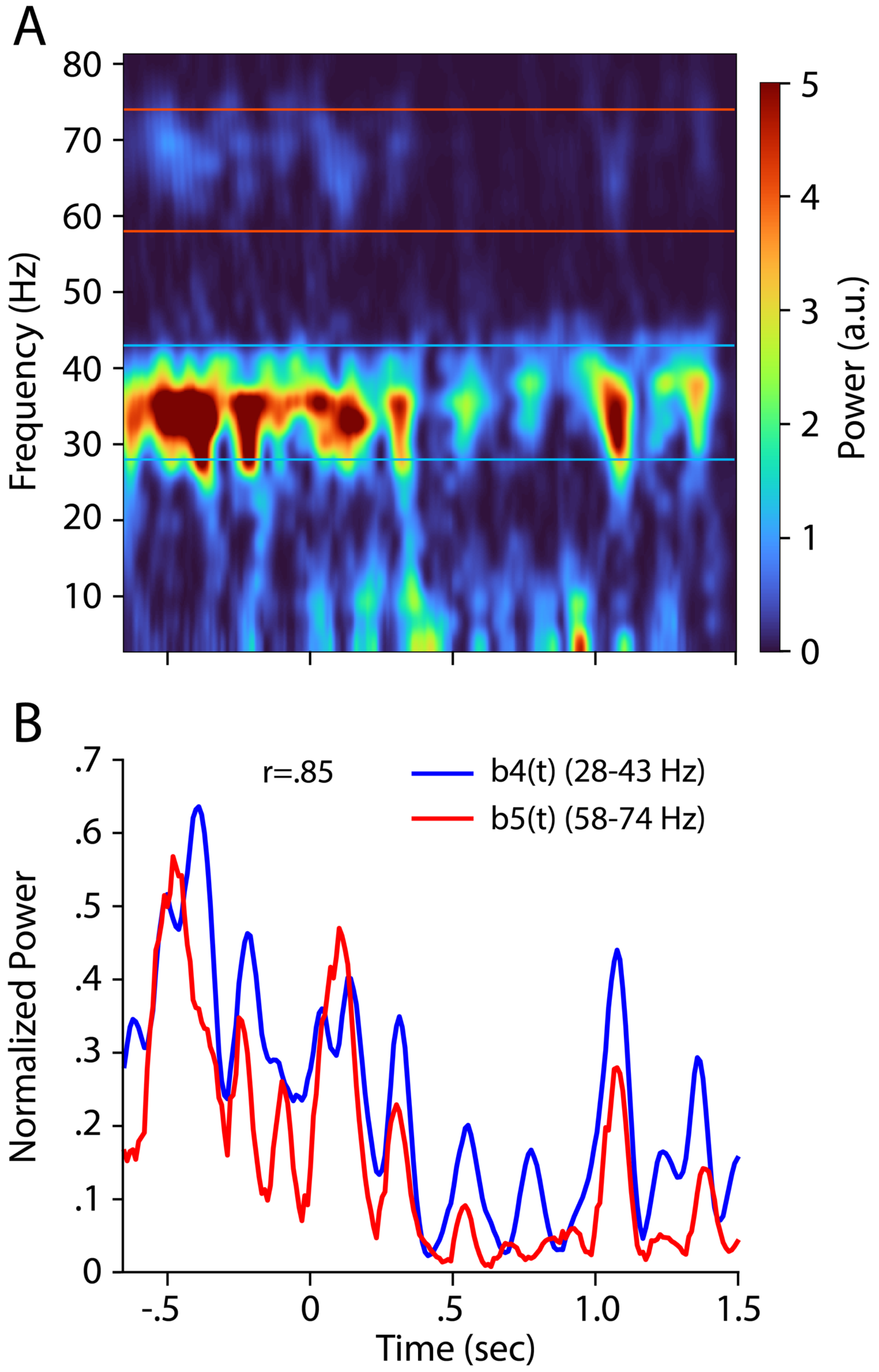

**Figure S4.** Example of the temporal correlation of band-4 and band-5 spectral envelopes b4(t) and b5(t). (A) Time-frequency spectrum of a single trial in area a3 of Monkey L. This channel displayed peaks in both band-4 and band-5 in the average spectrum (see Figure 1C, area a3). (B) Spectral envelopes b4(t) and b5(t) (normalized to the session maximum in each band) on the same trial. The blue and red lines in A bracket the frequencies used to compute b4(t) and b5(t) shown in B. The CC between the two signals is .85.

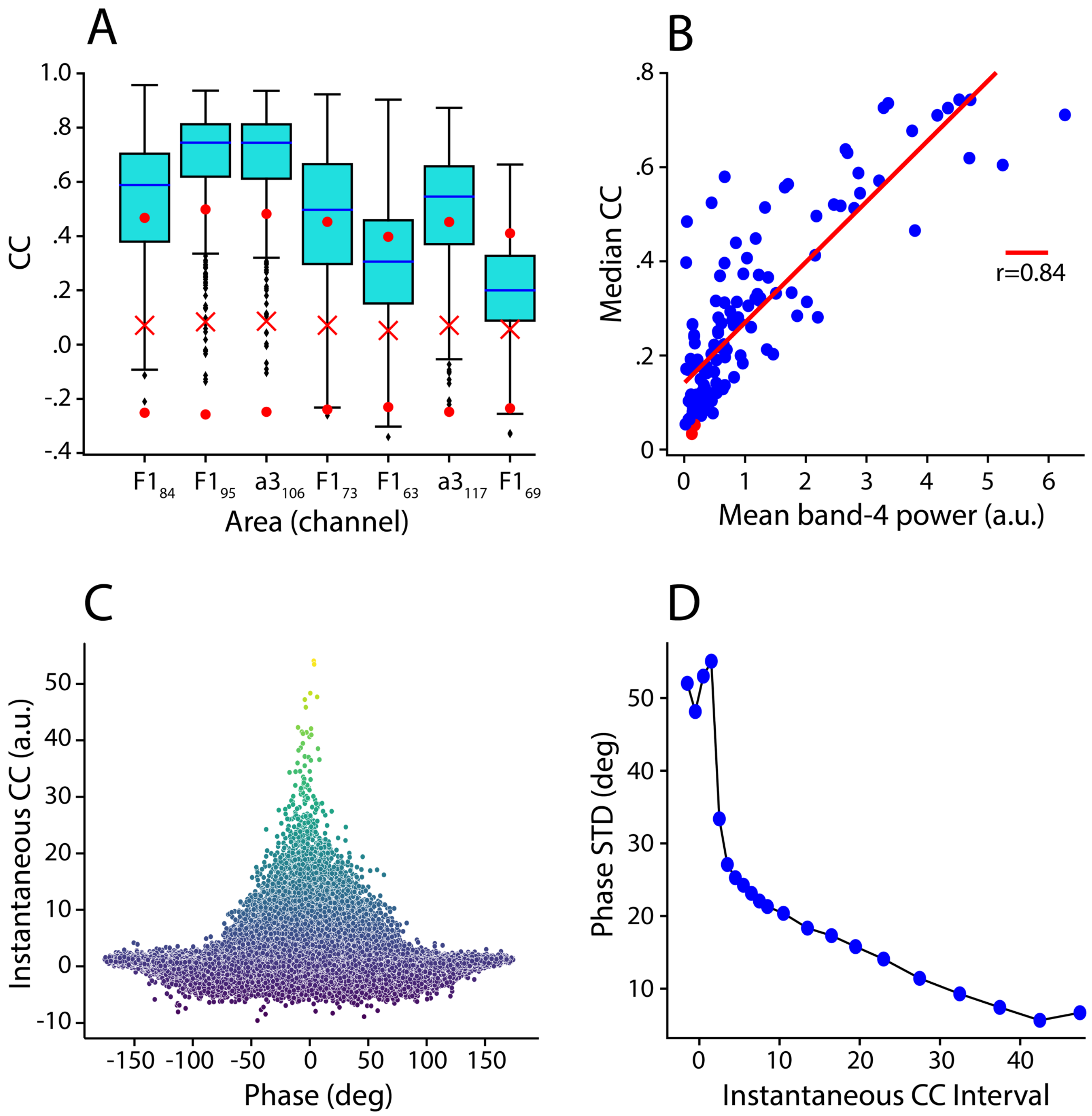

**Figure S5.** Cumulative results of correlation analysis between b4(t) and b5(t) in Monkey L. (A) Box plots of distributions of correlation coefficients (CC) across all trials on a single session for the channels containing spectral peaks in band-4 and band-5. The plots in Figure S4 are taken from channel 106, area a3. Red Xs and filled circles show the median and 95$^{th}$ and 5$^{th}$ percentiles for the randomized surrogate CC distributions. The observed distributions exceeded the surrogate distributions on all selected channels in this session (KS-test, p<<.001). (B) Scatter plot of the median CC vs the mean power in band-4 for all selected channels in the data set (n=132). The correlation coefficient between $b_4(t)$ and $b_5(t)$ increases with the amplitude of the band-4 component.

Blue data points indicate a significant difference from the corresponding surrogate distribution (KS-test, $p<.001$). (C) Scatter plot of the instantaneous correlation magnitude vs instantaneous phase between $b_4(t)$ and $b_5(t)$ for all the selected channels in the data set (n=132). (D) Standard deviation of the instantaneous phase distribution for discrete intervals of the instantaneous correlation magnitude, for all sessions, channels, trials and times. The phase difference between $b_4(t)$ and $b_5(t)$ decreases sharply as the correlation between the signals increases.

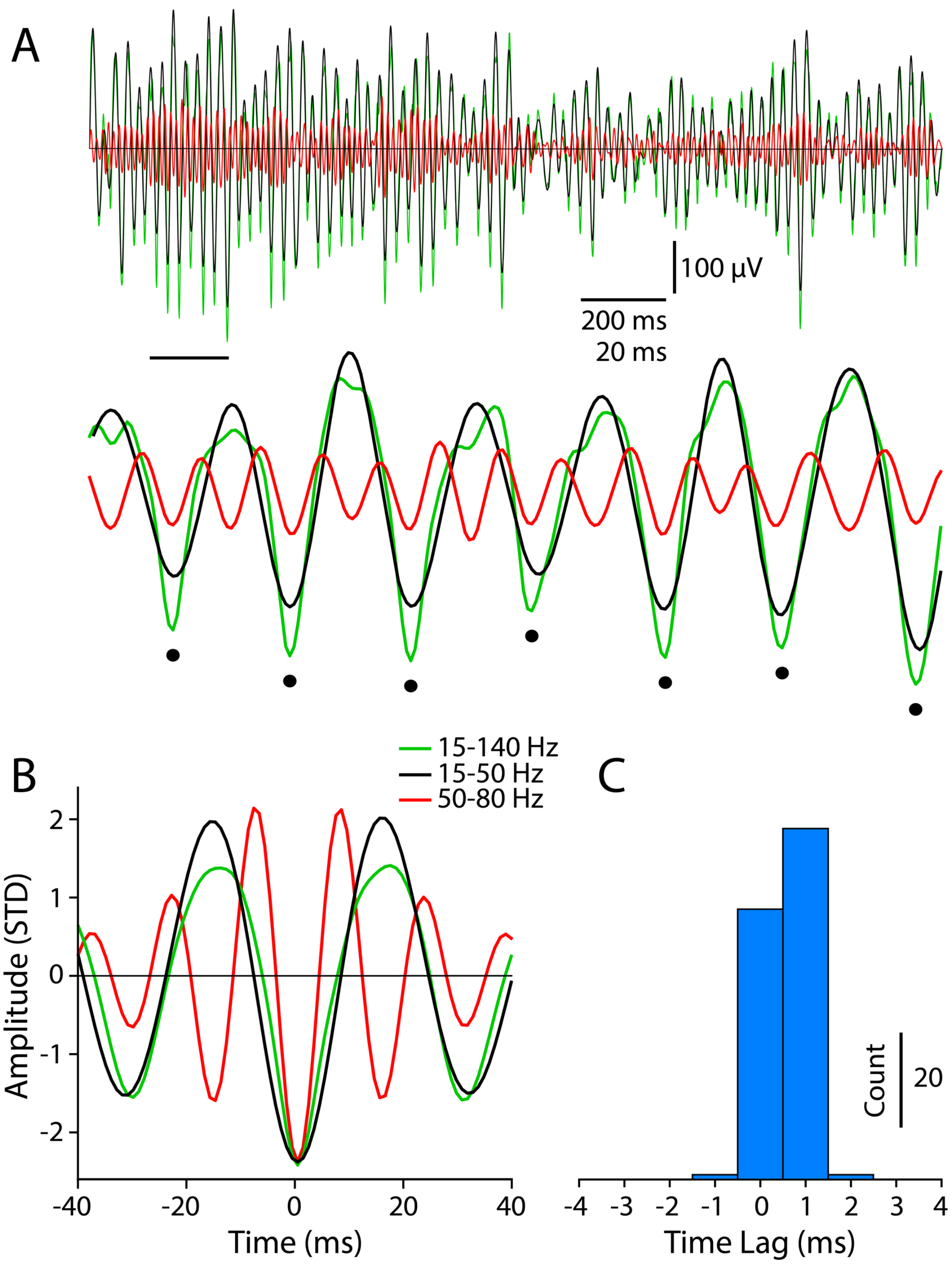

**Figure S6.** Non-sinusoidal shape of the band-4 oscillations and phase relationships between band-4 and band-5. (A) LFP data on the same channel and trial as shown in figure S4, filtered in 3 bands (blue: broadband (15-140 Hz); black: lowpass (15-50 Hz); red: highpass (50-80 Hz)). Black filled circles in the middle plot mark the time of negative peaks in the broadband (15-140 Hz) signal. The middle plot is from the data marked by the black horizontal line in the upper plot. Note the non-sinusoidal shape of the broadband oscillations and the tight phase coupling with the band-5 oscillations. (B) Cycle-triggered-averages (CTA) computed from the z-score normalized LFP on the same channel as shown in A. Averaging was triggered in the negative peaks of the broadband signal in the amplitude range of 2-3 standard deviations. The broadband CTA (blue) is noticeably more non-sinusoidal than the lowpass CTA (black). The negative peak of the highpass CTA (red) occurs near 0 ms, indicating that the band-5 oscillations are tightly phase locked to the peaks of the band-4 oscillations. (C) Histogram of the time lags of the highpass CTA for all channels containing spectral peaks in band-4 and band-5 (n=132). The narrow distribution centered at 0-1 ms demonstrates close phase locking between the band-4 and band-5 spectral components.

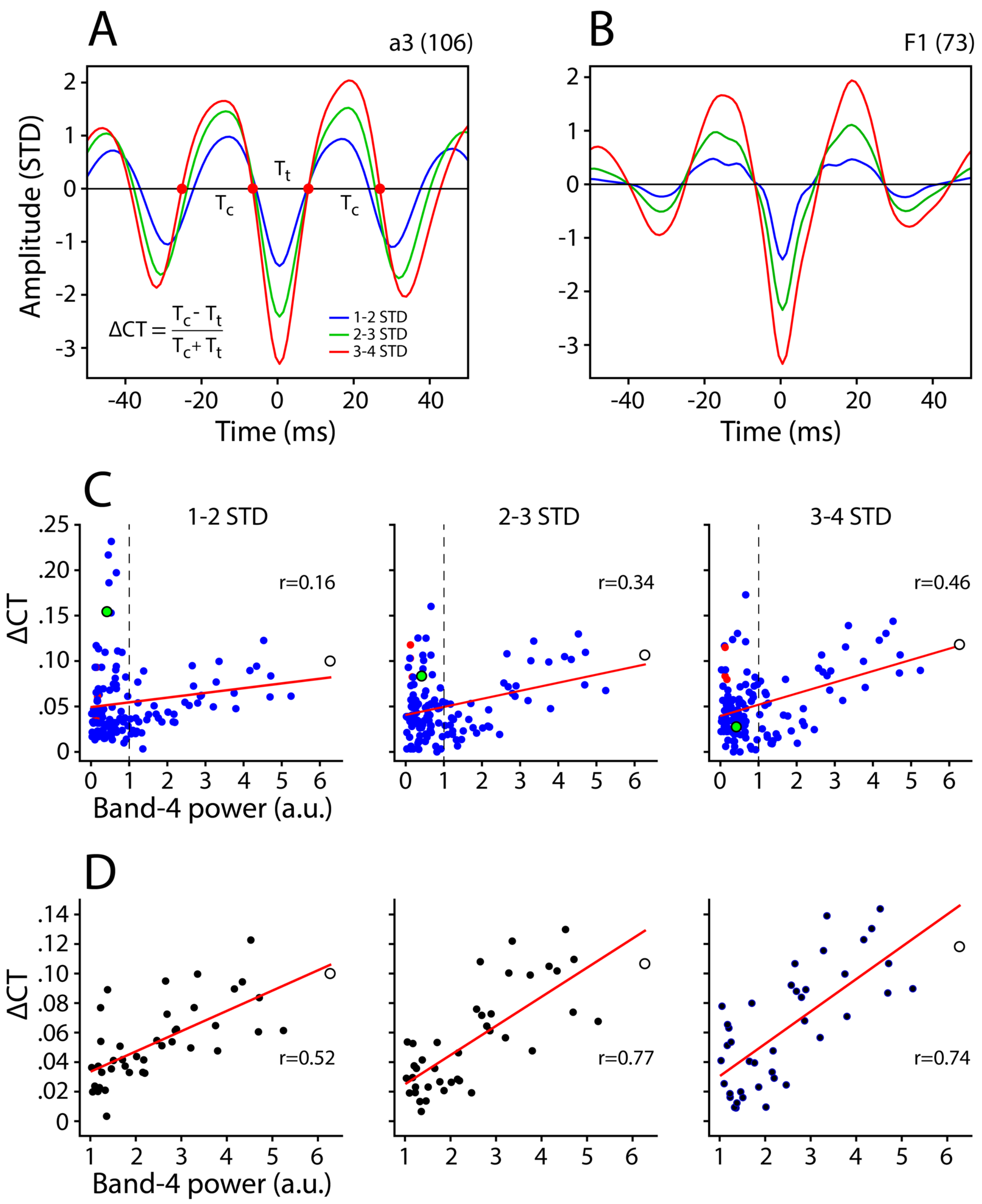

**Figure S7.** Waveform asymmetry (ΔCT) of the CTAs computed from the highpass signals and its relation to the amplitude of the band-4 power. (A,B) CTAs of the broadband signal (15-140 Hz) for two channels (a3, F1) on the same session shown in Figures S4, S5A and S6, computed at three different amplitude ranges (1-2, 2-3 and 3-4 stdevs). (C) Scatter plot of ΔCT vs band-4 power for all channels containing spectral peaks in band-4 and band-5 (n=132). The unfilled black circles and filled green circles show the values derived from the examples shown in A and B, respectively. (D) Same data as shown in C after removing values where band-4 power <1.

**Feature Importance**

We sought to determine which features in the classifier were responsible for successful classification. We first assessed the change in validation accuracy that occurs when the values of each feature are separately randomized. In this analysis the randomization of informative features should reduce validation accuracy (Dreyfus and Guyon, 2006). Figure S8 shows the results of this analysis applied to the presample epoch. There was a ~5% decrease in the mean validation accuracy across areas for each feature in both monkeys as compared to the baseline (band-3 SC and SampEn showed slightly larger reductions in both monkeys). This result suggests that each feature makes a small and approximately equal contribution to the validation accuracy.

However, given the possible redundancy between SC and PA, separately randomizing each feature doesn't assess the relative contribution of SC versus PA or the contribution of each frequency band. We therefore ran two additional decoding analyses after removing SampEn as a feature. We assessed the validation accuracy independently for SC and PA using all frequency bands (we refer to this as amplitude analysis) and we assessed the effect on validation accuracy of removing both features in each frequency band separately (we refer to this as frequency analysis). The results, shown in Tables S1 and S2, reveal mostly weak, but widespread effects in nearly all areas. For the amplitude analysis, removing SC or PA in all frequency bands resulted in a significant difference in validation accuracy for all 11 areas in monkey E and in 23 out of 28 areas in monkey L. For some areas SC was more informative than PA while the opposite effect was present in other areas. The only apparent trend occurred in monkey L where validation accuracy was greater for PA than SC in 16 out 23 areas. For the frequency analysis, significant differences in validation accuracy occurred in all 11 areas of monkey E and in 27 out of 28 areas in monkey L. The most informative frequency bands (i.e., those resulting in the greatest reduction of validation accuracy when omitted) were bands 2-4 of monkey E and bands 3-5 of monkey L. These effects were roughly evenly distributed across areas in both monkeys, supporting the notion that most features make a small contribution to the validation accuracy.

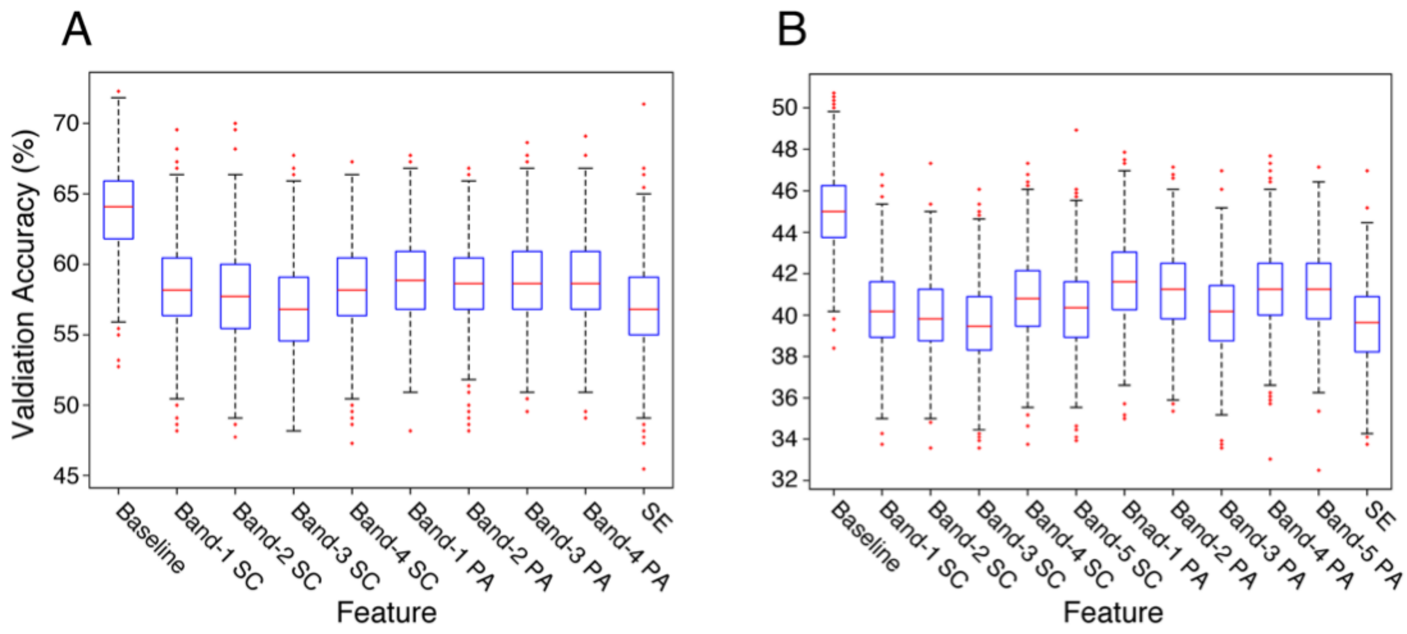

**Figure S8.** Contribution of features to validation accuracy. Box plots of the mean validation accuracy across all areas during the presample epoch when each feature is separately randomized. The plots in A and B show the results for monkeys E and L, respectively. The control distribution (baseline) is shown in the leftmost box of each plot. A drop of classification accuracy reflects the importance of that feature for correct classification.

**Table S1.** Mean validation accuracies for the listed features (SampEn excluded) during the presample epoch for monkey E. The baseline values, shown for reference, include all features in all bands (SC, PA, SampEn). Asterisks indicate significant differences ($p<10^{-5}$). For the amplitude analysis the asterisk denotes the greater of the two values (Mann Whitney U-test, FDR corrected). For the frequency analysis the asterisk denotes the smallest value (Kruskal-Wallis test, FDR corrected). FDR is corrected for number of areas (n=11).

| Monkey E | | | Amplitude Analysis | | | Frequency Analysis | | | |
|---|---|---|---|---|---|---|---|---|---|
| Areas | Baseline | | Just SC | Just PA | | Omit | Omit | Omit | Omit |
| dPFC | 81.87 | | 83.46* | 74.40 | | 81.07 | 80.11 | 71.29* | 79.97 |
| F1 | 49.63 | | 27.48 | 49.87* | | 45.86 | 44.21 | 42.51* | 43.13 |
| F6/7 | 77.80 | | 72.42 | 77.91* | | 74.71 | 77.08 | 69.01* | 78.38 |
| a2 | 76.87 | | 65.87* | 59.49 | | 70.18* | 72.22 | 73.67 | 70.53 |
| 7b | 76.64 | | 77.69* | 54.96 | | 75.63 | 73.69* | 75.66 | 78.95 |
| DP | 37.90 | | 37.36* | 30.75 | | 31.65 | 29.25 | 34.95 | 27.64* |
| V6A | 50.90 | | 57.18* | 39.85 | | 55.36 | 43.62 | 47.21 | 42.16* |
| V4/t | 51.64 | | 40.03 | 46.32* | | 49.59 | 51.05 | 49.80 | 45.06* |
| V2 | 59.69 | | 54.65* | 47.98 | | 55.59 | 52.76 | 54.70 | 52.01* |
| $V1_{SR}$ | 74.37 | | 64.37 | 69.82* | | 73.79 | 68.94* | 72.03 | 73.01 |
| V1 | 65.08 | | 42.68 | 59.24* | | 62.40 | 58.81* | 59.43 | 61.49 |
| Mean | 63.85 | | 56.65 | 55.51 | | 61.44 | 59.25 | 59.11 | 59.30 |

**Table S2.** Mean validation accuracies for the listed features (SampEn excluded) during the presample epoch for monkey L. The baseline values, shown for reference, include all features in all bands (SC, PA, SampEn). Asterisks indicate significant differences ($p<10^{-5}$). For the amplitude analysis the asterisk denotes the greater of the two values (Mann Whitney U-test, FDR corrected). For the frequency analysis the asterisk denotes the smallest value (Kruskal-Wallis test, FDR corrected). FDR is corrected for number of areas (n=28).

| Monkey L | | Amplitude Analysis (all frequency bands) | | Frequency Analysis (all amplitudes) | | | | |
|---|---|---|---|---|---|---|---|---|
| Areas | Baseline | Just SC | Just PA | Omit | Omit | Omit | Omit | Omit |
| OrbPFC | 44.84 | 39.73 | 43.33* | 46.02 | 44.01 | 44.72 | 40.15* | 45.36 |
| 24 | 60.03 | 42.66 | 49.70* | 61.81 | 61.15 | 55.36* | 58.08 | 58.48 |
| vPFC | 38.47 | 21.25 | 27.49* | 37.19 | 38.49 | 35.95 | 32.90* | 36.23 |
| dPFC | 50.74 | 31.64 | 43.08* | 46.39 | 52.97 | 39.83 | 44.76 | 39.77* |
| 8B | 50.84 | 35.57 | 44.35* | 52.11 | 48.28 | 46.39* | 49.28 | 48.67 |
| 8L | 66.70 | 42.47 | 50.87* | 60.05 | 58.96 | 58.63 | 52.25* | 56.01 |
| 8M | 57.02 | 39.23 | 49.06* | 57.21 | 52.30* | 53.27 | 52.55 | 57.03 |
| 8r | 70.97 | 39.56 | 67.54* | 73.81 | 74.98 | 63.24* | 65.42 | 64.44 |
| 44/45 | 49.89 | 42.01* | 34.31 | 51.37 | 47.00 | 49.98 | 44.89* | 49.36 |
| F1 | 49.82 | 44.08 | 44.13 | 49.52 | 51.49 | 50.25 | 48.67 | 45.92* |
| F2 | 52.86 | 52.95* | 43.86 | 53.80 | 52.69 | 51.12 | 48.72 | 46.07* |
| F6/7 | 42.47 | 26.23 | 29.47* | 34.41 | 35.06 | 26.05* | 35.47 | 30.51 |
| a1 | 54.65 | 49.02 | 49.77 | 49.22* | 53.20 | 49.62 | 52.61 | 56.14 |
| a2 | 46.65 | 55.32* | 35.26 | 45.52 | 46.83 | 42.87* | 45.01 | 45.66 |
| a3 | 34.85 | 24.23 | 32.25* | 32.37 | 34.36 | 33.21 | 33.08 | 30.41* |
| 5/MIP | 34.47 | 19.58 | 28.60* | 31.75 | 30.93 | 29.87* | 32.60 | 32.38 |
| PIP | 53.75 | 37.92 | 52.07* | 55.29 | 52.66 | 52.91 | 51.18* | 56.75 |
| 7a | 20.71 | 10.42 | 17.62* | 17.62 | 17.47* | 18.89 | 19.06 | 18.68 |
| AIP/VIP | 38.38 | 34.28 | 35.31 | 37.84 | 33.72 | 31.88* | 38.41 | 34.40 |
| 7b | 56.19 | 38.70 | 59.19* | 53.23 | 53.06 | 51.67 | 58.30 | 45.63* |
| [illegible] | [illegible] | [illegible] | [illegible] | [illegible] | [illegible] | [illegible] | [illegible] | [illegible] |
| DP | 28.98 | 31.48* | 24.06 | 27.57 | 25.84 | 26.24 | 25.47 | 25.56 |
| V6A | 44.82 | 40.80 | 40.73 | 45.71 | 44.70 | 41.23 | 38.63* | 41.01 |

| | | | | | | | | | | |
|---|---|---|---|---|---|---|---|---|---|---|
| MT/MST | 22.09 | | 30.98* | 15.87 | | 20.75 | 22.50 | 21.42 | 19.52* | 22.48 |
| V4/t | 42.46 | | 27.67 | 36.18* | | 40.58 | 40.39 | 39.83 | 41.03 | 36.07* |
| V2$_{SR}$ | 35.72 | | 34.54* | 32.37 | | 33.91 | 33.78 | 34.66 | 32.86* | 37.53 |
| V2 | 36.95 | | 37.12* | 32.09 | | 34.15 | 31.50* | 33.97 | 37.00 | 37.54 |
| V1 | 51.21 | | 47.52 | 48.71 | | 51.26 | 51.01 | 48.27* | 50.86 | 50.76 |
| Mean | 45.07 | | 35.36 | 39.04 | | 43.79 | 43.23 | 41.11 | 41.84 | 41.89 |

**Areal Misclassification**

What is responsible for the wide range of validation accuracies across cortical areas? We considered two interrelated factors that might introduce confusion in the classification analysis. First, the spectral and temporal features of the LFP could vary widely within an area, making the data difficult to separate. Second, as indicated by the flatmaps of SC, there could be a high degree of feature similarity between areas, particularly if they are close to one another. We examined the first question by calculating the correlation coefficient between validation accuracy and the standard deviation of each feature used in the classifier (Table S3). Significant negative correlations were present for 3 out of 9 features in monkey E, and 4 out of 11 features in monkey L. These effects occurred at lower frequencies, in bands 1 and 2, and no correlation was observed for SampEn.

To determine how validation accuracy depends on the misclassification between areas, we calculated a "pairwise confusability" metric from the confusion matrices. This metric is the sum of the off-diagonal classification errors between each pair of areas in the confusion matrix. The resulting values were used to create a similarity matrix where all pairwise combinations of areas are represented, and each value indicates the percentage of incorrect classifications between each pair. We used this similarity matrix as input to a multi-dimensional scaling algorithm to visualize the off-diagonal misclassifications in the confusion matrix (Figure S9). In these plots, the distance between points represents the magnitude of separation among the spectral and temporal features of all the areas. Interestingly, the data from both monkeys appeared to segregate into regional clusters (primary visual, extrastriate visual and fronto-parietal in Monkey E, and occipito-parietal, somatomotor and premotor-prefrontal in Monkey L). This provides further evidence that the spectral and temporal features of the LFP exhibit spatial gradients with nearby areas displaying similar profiles.

**Table S3.** Correlation coefficients, and corresponding P values, between the standard deviation of classification features and validation accuracy for the Presample epoch of the task. SC – Spectral Content; PA – Peak Amplitude, SampEn – Sample Entropy. Numerical values 1-5 indicate the separate frequency bands for each monkey.

| **Monkey E** | SC1 | SC2 | SC3 | SC4 | | PA1 | PA2 | PA3 | PA4 | | SampEn |
|---|---|---|---|---|---|---|---|---|---|---|---|
| Correlation | -.75 | -.66 | -.28 | .28 | | -.84 | -.16 | -.44 | -.54 | | -.05 |
| P value | .008 | .029 | .41 | .41 | | .001 | .63 | .18 | .09 | | .89 |
| **Monkey L** | SC1 | SC2 | SC3 | SC4 | SC5 | PA1 | PA2 | PA3 | PA4 | PA5 | SampEn |
| Correlation | -.48 | -.45 | .16 | .02 | .14 | -.56 | -.61 | -.35 | -.06 | -.20 | -.11 |
| P value | .01 | .017 | .42 | .91 | .47 | .002 | .0006 | .07 | .75 | .30 | .59 |

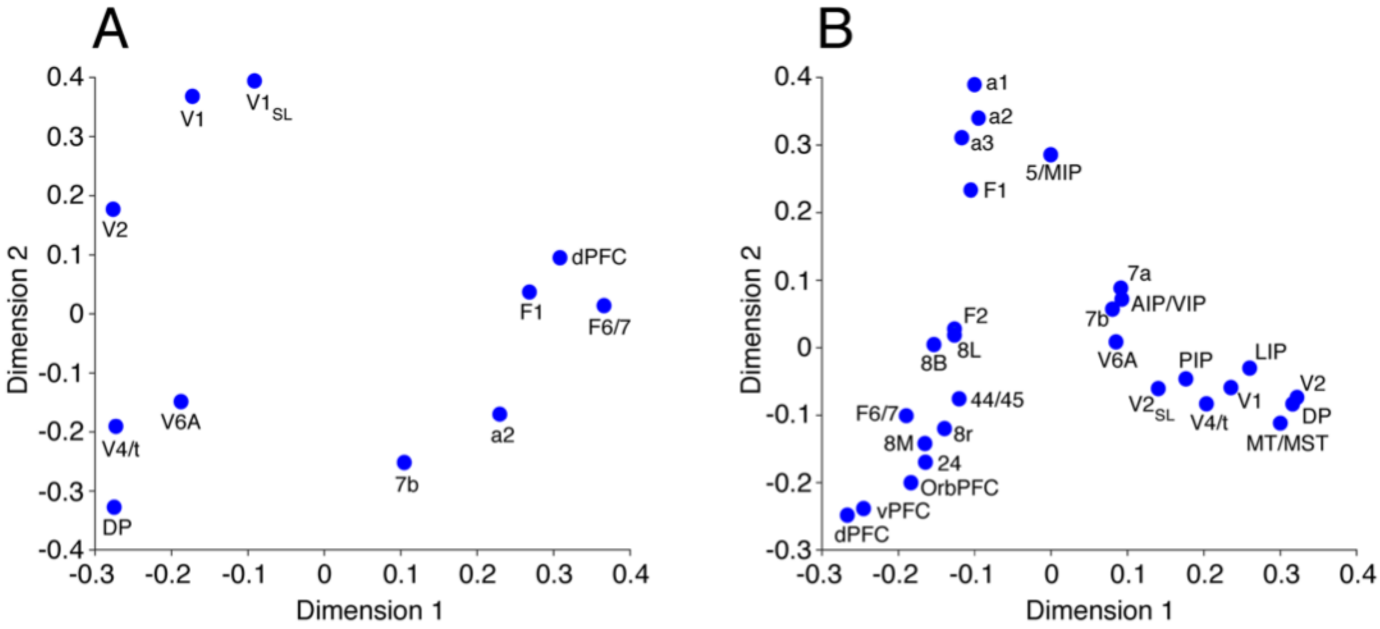

**Figure S9.** Multidimensional scaling maps of the pairwise confusability in monkey E (A) and monkey L (B). The distance between points represents the magnitude of separation among the spectral and temporal features of all the area/groups.
